# The *R2R3-MYB* gene family in banana (*Musa acuminata*): genome-wide identification, classification and expression patterns

**DOI:** 10.1101/2020.02.03.932046

**Authors:** Boas Pucker, Ashutosh Pandey, Bernd Weisshaar, Ralf Stracke

## Abstract

The *R2R3-MYB* genes comprise one of the largest transcription factor gene families in plants, playing regulatory roles in plant-specific developmental processes, defense responses and metabolite accumulation. To date MYB family genes have not yet been comprehensively identified in the major staple fruit crop banana. In this study, we present a comprehensive, genome-wide analysis of the *MYB* genes from *Musa acuminata* DH-Pahang (A genome). A total of 285 *R2R3-MYB* genes as well as genes encoding three other classes of MYB proteins containing multiple MYB repeats were identified and characterised with respect to structure and chromosomal organisation. Organ- and development-specific expression patterns were determined from RNA-seq data. For 280 *M. acuminata MYB* genes for which expression was found in at least one of the analysed samples, a variety of expression patterns were detected. The *M. acuminata R2R3-MYB* genes were functionally categorised, leading to the identification of seven clades containing only *M. acuminata* R2R3-MYBs. The encoded proteins may have specialised functions that were acquired or expanded in *Musa* during genome evolution. This functional classification and expression analysis of the *MYB* gene family in banana establishes a solid foundation for future comprehensive functional analysis of MaMYBs and can be utilized in banana improvement programmes.

## Introduction

Banana (*Musa* spp.), including dessert and cooking types, is a staple fruit crop for a major world population, especially in developing countries. The crop is grown in more than 100 countries throughout the tropics and sub-tropics, mainly in the African, Asia-Pacific, and Latin American and Caribbean regions [1]. Bananas provide an excellent source of energy and are rich in certain minerals and in vitamins A, C and B6. Furthermore, this perennial, monocotyledonous plant provides an important source of fibre, sugar, starch and cellulose (used for paper, textiles). Bananas have also been considered as a useful tool to deliver edible vaccines [2]. Certain agronomic traits, such as stress and pest resistance as well as fruit quality, are thus of considerable interest. Banana improvement through breeding exercises has been challenging for various reasons. Therefore, genetic engineering-based optimisations hold great promise for crop improvement. For this purpose, candidate gene targets need to be identified. The release of a high quality banana genome sequence [3] provides an useful resource to understand functional genomics of important agronomic traits and to identify candidate genes to be utilized in banana improvement programmes.

Almost all biological processes in eukaryotic cells or organisms are influenced by transcriptional control of gene expression. Thus, the regulatory level is a good starting point for genetic engineering. Regulatory proteins are involved in transcriptional control, alone or complexed with other proteins, by activating or repressing (or both) the recruitment of RNA polymerase to promoters of specific genes [4]. These proteins are called transcription factors. As expected from their substantial regulatory complexity, transcription factors are numerous and diverse [5]. By binding to specific DNA sequence motifs and regulating gene expression, transcription factors control various regulatory and signaling networks involved in the development, growth and stress response in an organism.

One of the widest distributed transcription factor families in all eukaryotes is the MYB (myeloblastosis) protein family. In the plant kingdom, MYB proteins constitute one of the largest transcription factor families. MYB proteins are defined by a highly conserved MYB DNA-binding domain, mostly located at the N-terminus of the protein. The MYB domain generally consists of up to four imperfect amino acid sequence repeats (R) of about 50-53 amino acids, each forming three alpha–helices [summarised in 6]. The second and third helices of each repeat build a helix–turn–helix (HTH) structure with three regularly spaced tryptophan (or hydrophobic) residues, forming a hydrophobic core [7]. The third helix of each repeat was identified as the DNA recognition helix that makes direct contact with DNA [8].

During DNA contact, two MYB repeats are closely packed in the major groove, so that the two recognition helices bind cooperatively to the specific DNA recognition sequence motif. In contrast to vertebrates genomes, which only encode MYB transcription factors with three repeats, plants have different MYB domain organisations, comprising one to four repeats [6, 9]. R2R3-MYBs, which are MYB proteins with two repeats (named according to repeat numbering in vertebrate MYBs), are particularly expanded in plant genomes. Copy numbers range from 45 unique *R2R3-MYBs* in *Ginkgo biloba* [10] to 360 in Mexican cotton (*Gossypium hirsutum*) [11]. The expansion of the R2R3-MYB family was coupled with widening in the functional diversity of R2R3-MYBs, considered to regulate mainly plant-specific processes including secondary metabolism, stress responses and development [6]. As expected, R2R3-MYBs are involved in regulating several biological traits, for example wine quality, fruit color, cotton fibre length, pollinator preferences and nodulation in legumes.

In this study, we have used genomic resources to systematically identify members of the *M. acuminata* (A genome) *R2R3-MYB* gene family. We used knowledge from other plant species, including the model plant *A. thaliana*, leading to a functional classification of the banana *R2R3-MYB* genes based on the MYB phylogeny. Furthermore, RNA-seq data was used to analyse expression in different *M. acuminata* organs and developmental stages and to compare expression patterns of closely grouped co-orthologs. The identification and functional characterization of the *R2R3-MYB* gene family from banana will provide an insight into the regulatory aspects of different biochemical and physiological processes, as those operating during fruit ripening as well as response to various environmental stresses.Our findings offer the first step towards further investigations on the biological and molecular functions of MYB transcription factors with the selection of genes responsible for economically important traits in *Musa*, which can be utilized in banana improvement programmes.

## Material and Methods

### Search for MYB protein coding genes in the *M. acuminata* genome

A consensus R2R3-MYB DNA binding domain sequence [12] (S1 Table) was used as protein query in tBLASTn [13] searches on the *M. acuminata* DH-Pahang genome sequence (version 2) (https://banana-genome-hub.southgreen.fr/sites/banana-genome-hub.southgreen.fr/files/data/fasta/version2/musa_acuminata_v2_pseudochromosome.fna) in an initial search for MYB protein coding genes. To confirm the obtained amino acid sequences, the putative MYB sequences were manually analysed for the presence of an intact MYB domain. All *M. acuminata* MYB candidates from the initial BLAST were inspected to ensure that the putative gene models encode two or more (multiple) MYB repeats. The identified gene models were analysed to map them individually to unique loci in the genome and redundant sequences were discarded from the data set to obtain unique *MaMYB* genes.

The identified *MaMYB* genes were matched to the automatically annotated genes from the Banana Genome Hub database [14]. The open reading frames of the identified *MaMYB* were manually inspected and, if available, verified by mapping of RNA-seq data to the genomic sequence. Resulting *MaMYB* annotation is provided in the supporting information: Multi-FASTA files with *MaMYB* CDS sequences (S1 File) and MaMYB peptide sequences (S2 File), general feature format (GFF) file (S3 File) describing the *MaMYB* genes for use in genome viewers/browers.

### Genomic distribution of *MaMYB* genes

The *MaMYB* genes were located on the corresponding chromosomes by the MapGene2chromosome web v2 (MG2C) software tool (http://mg2c.iask.in/mg2c_v2.0/) according to their position information of a physical map, available from the DH-Pahang genome annotation (V2).

### Phylogenetic analyses

Protein sequences of 133 *A. thaliana* MYBs were obtained from TAIR (http://www.arabidopsis.org/). We also considered other multiple MYB-repeat proteins from *A. thaliana* in the phylogenetic analysis to determine orthologs in the *M. acuminata* genome: five AtMYB3R, AtMYB4R1 and AtCDC5. Additionally, 43 well-known, functionally characterised landmark plant R2R3-MYB protein sequences were collected from GenBank at the National Center for Biotechnology Information (NCBI) (http://www.ncbi.nlm.nih.gov/). Phylogenetic trees were constructed from ClustalΩ [15] aligned MYB domain sequences (293 MaMYBs, 132 AtMYBs and 43 plant landmark MYBs) using MEGA7 [16] with default settings. A mojority rule Maximum Likelihood (ML) consensus tree inferred from 1000 bootstrap replications was calculated. *M. acuminata* MYB proteins were classified according to their relationships with corresponding *A. thaliana* and landmark MYB proteins.

### Expression analysis from RNA-seq data

RNA-Seq data sets were retrieved from the Sequence Read Archive (https://www.ncbi.nlm.nih.gov/sra) via fastq-dump v2.9.6 (https://github.com/ncbi/sra-tools) (S2 Table). STAR v2.5.1b [17] was applied for the mapping of reads to the Pahang v2 reference genome sequence [18] using previously described parameters [19]. featureCounts [20] was applied for quantification of the mapped reads per gene based on the Pahang v2 annotation. Previously developed Python scripts were deployed for the calculation of normalized expression values (https://github.com/bpucker/bananaMYB) [19].

## Results and Discussion

The first annotated reference genome sequence of *M. acuminata* (A genome) became available in 2012 [3]. It was obtained from a double haploid (DH) plant of the Pahang cultivar, derived through haploid pollen and spontaneous chromosome doubling from the wild subspecies *Musa acuminata* ssp. *malaccensis* [3]. This wild subspecies was involved in the domestication of the vast majority of cultivated bananas and its genetic signature is commonly found in dessert and cooking bananas (ProMusa, http://www.promusa.org). An improved version (DH-Pahang version 2) of the genome assembly and annotation was presented in 2016 comprising 450.7 Mb (86% of the estimated size) from which 89.5% are assigned to one of the 11 chromosomes, predicted to contain 35,276 protein encoding genes [18]. This *M. acuminata* DH-Pahang version 2 genome sequence provides the platform of this study.

### Identification and genomic distribution of *M. acuminata MYB* genes

A consensus R2R3-MYB DNA binding domain sequence (deduced from Arabidopsis, grape and sugarbeet R2R3-MYBs, S1 Table) was used as protein query in tBLASTn searches on the DH-Pahang version 2 genome sequence to comprehensively identify MYB protein coding genes in *M. acuminata*. The resulting putative MYB sequences were proven to map to unique loci in the genome and were confirmed to contain an intact MYB domain. This ensured that the gene models contained two or more (multiple) MYB repeats. We identified a set of 285 R2R3-MYB proteins and nine multiple repeat MYB proteins distantly related to the typical R2R3-MYB proteins: six R1R2R3-MYB (MYB3R) proteins, two MYB4R proteins and one CDC5-like protein from the *M. acuminata* genome sequence (Table 1).

**Table 1.**
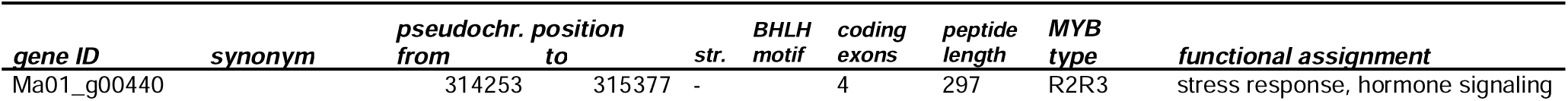
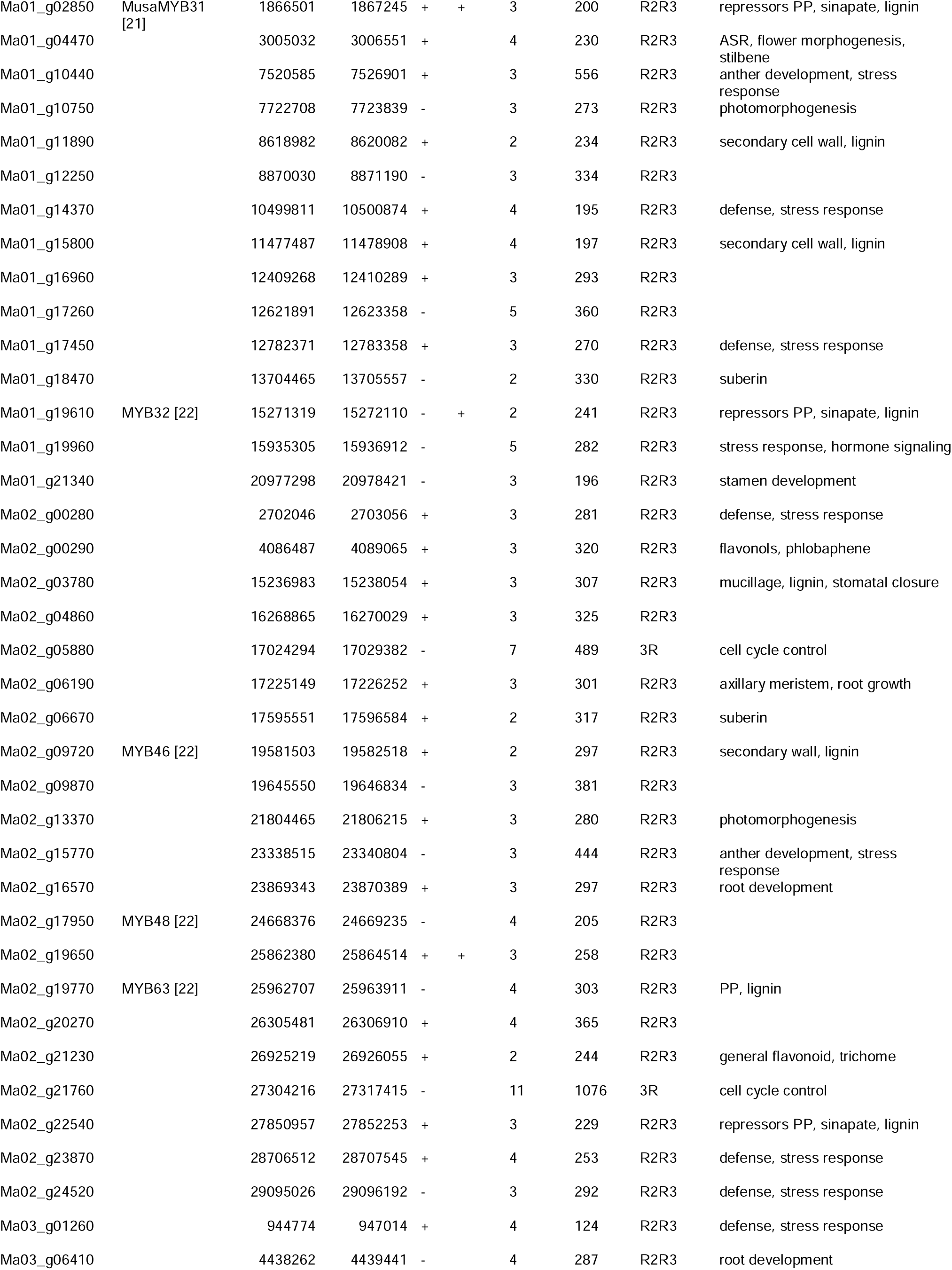
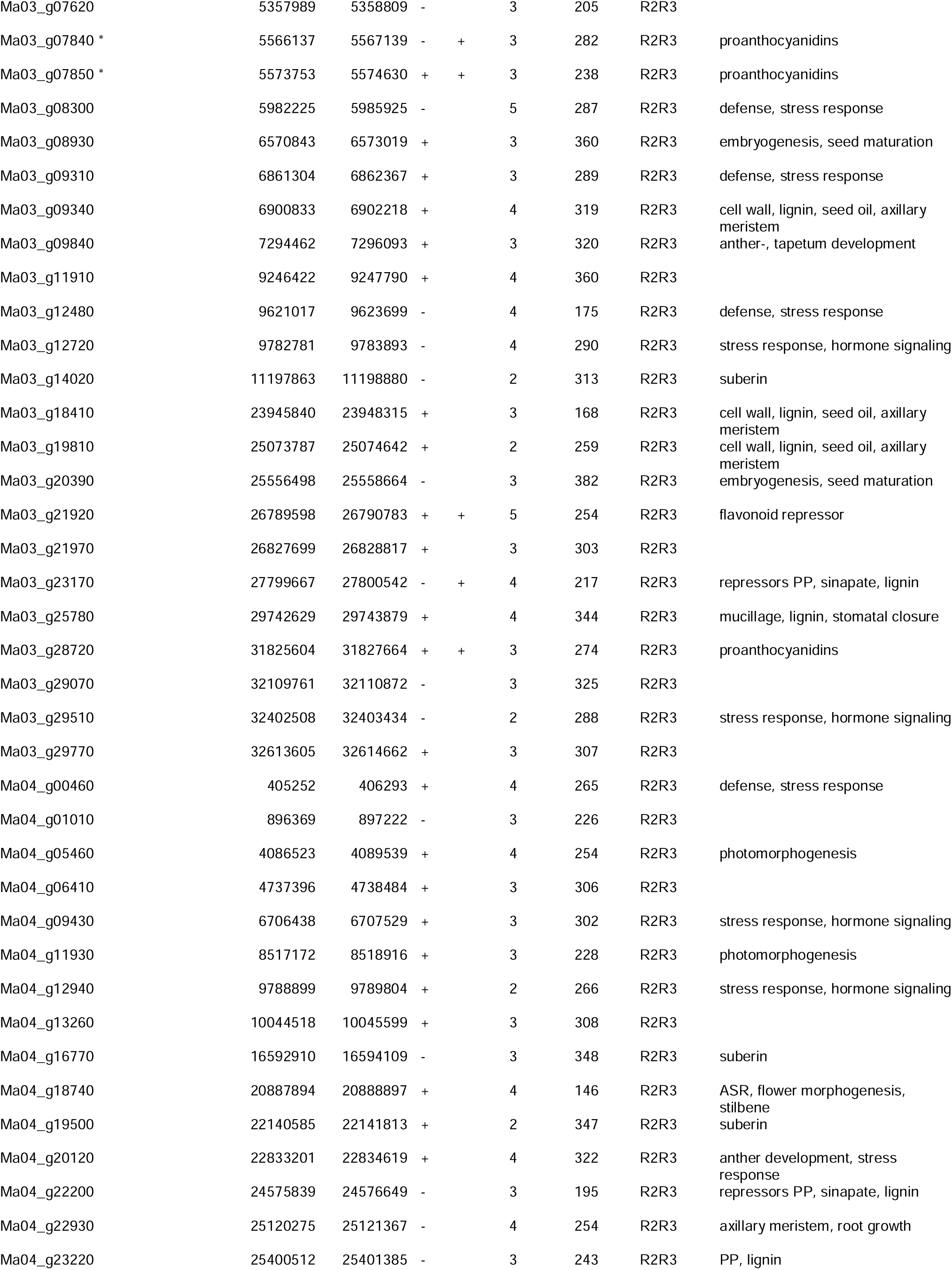
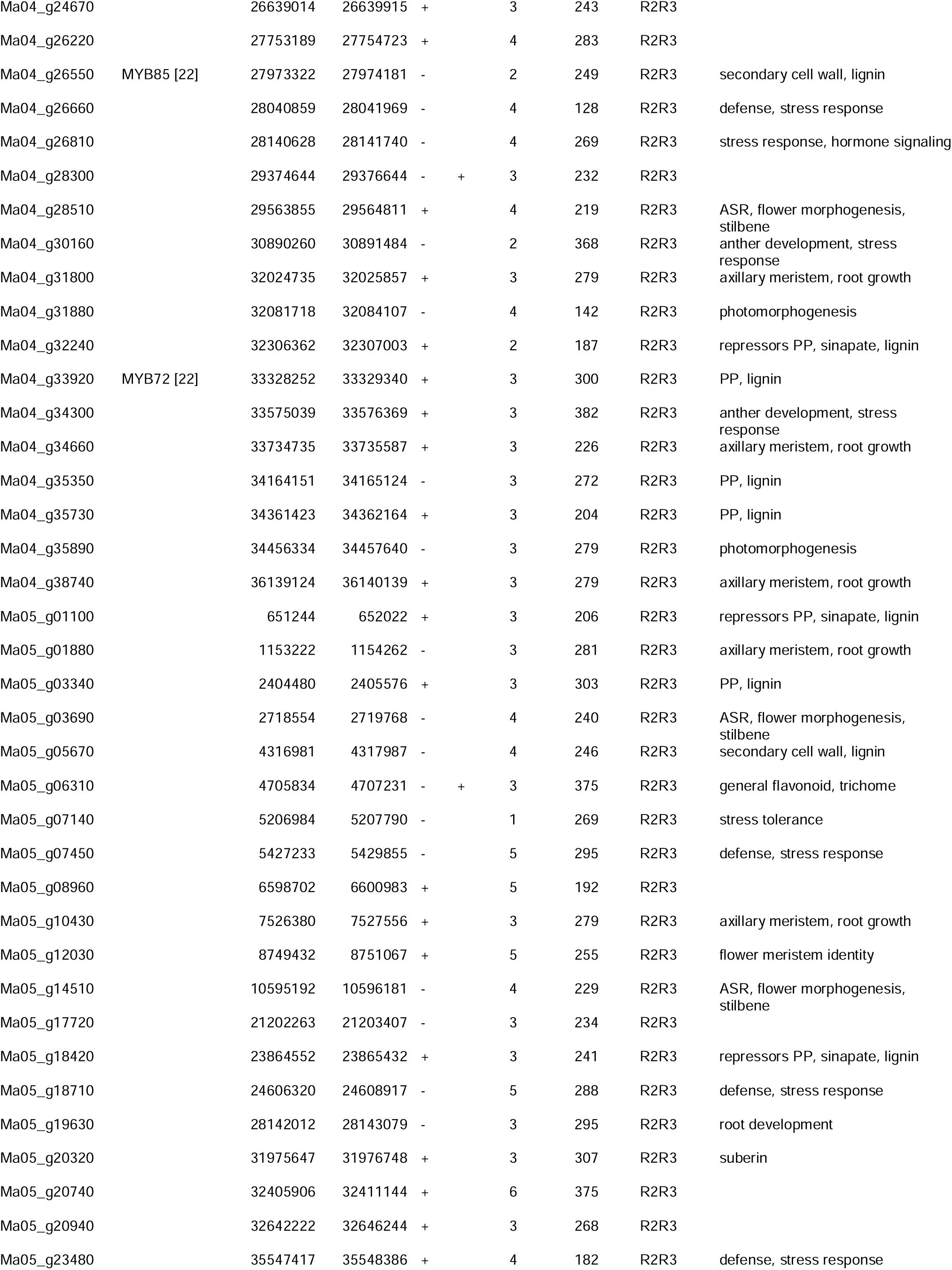
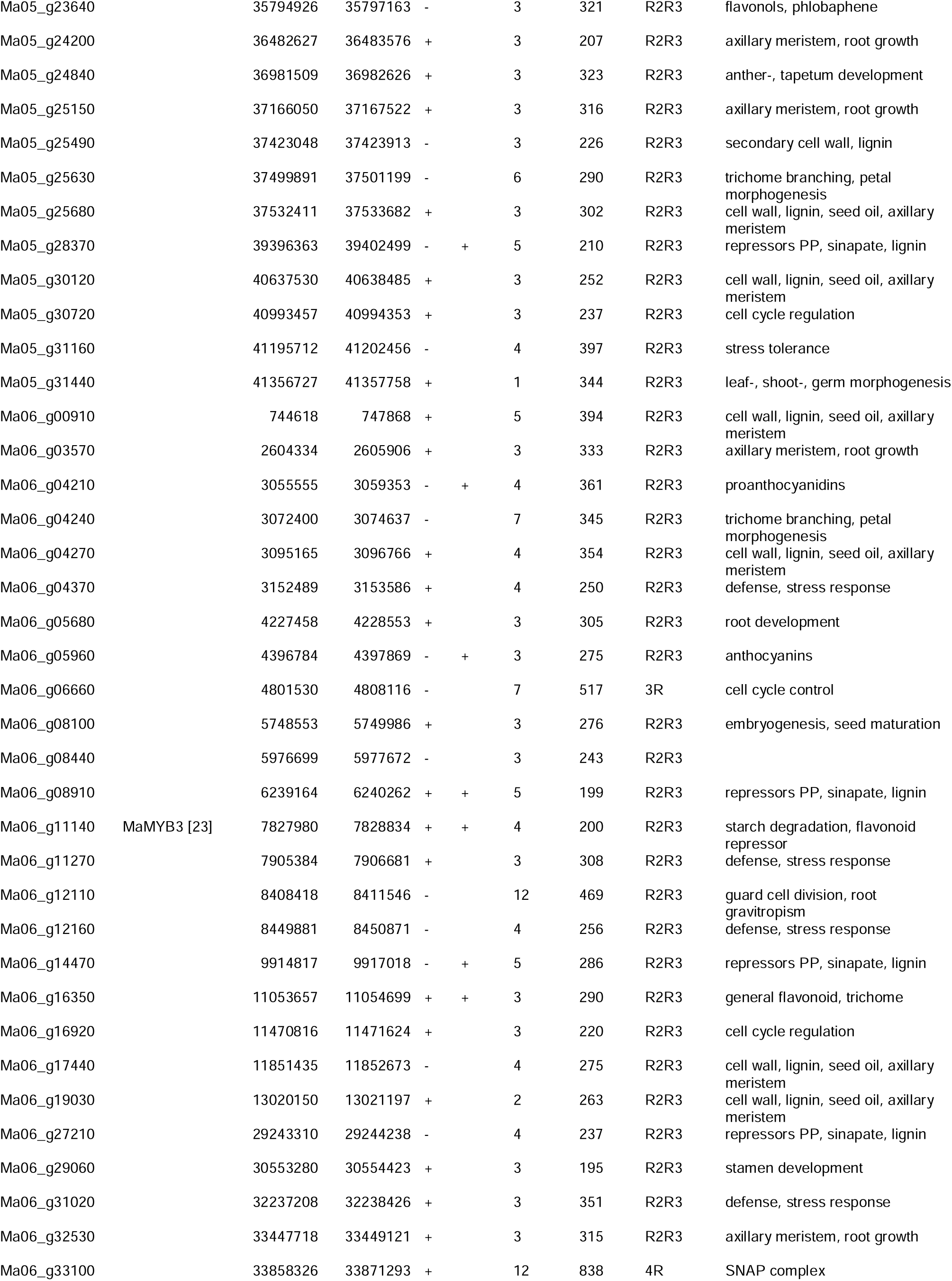
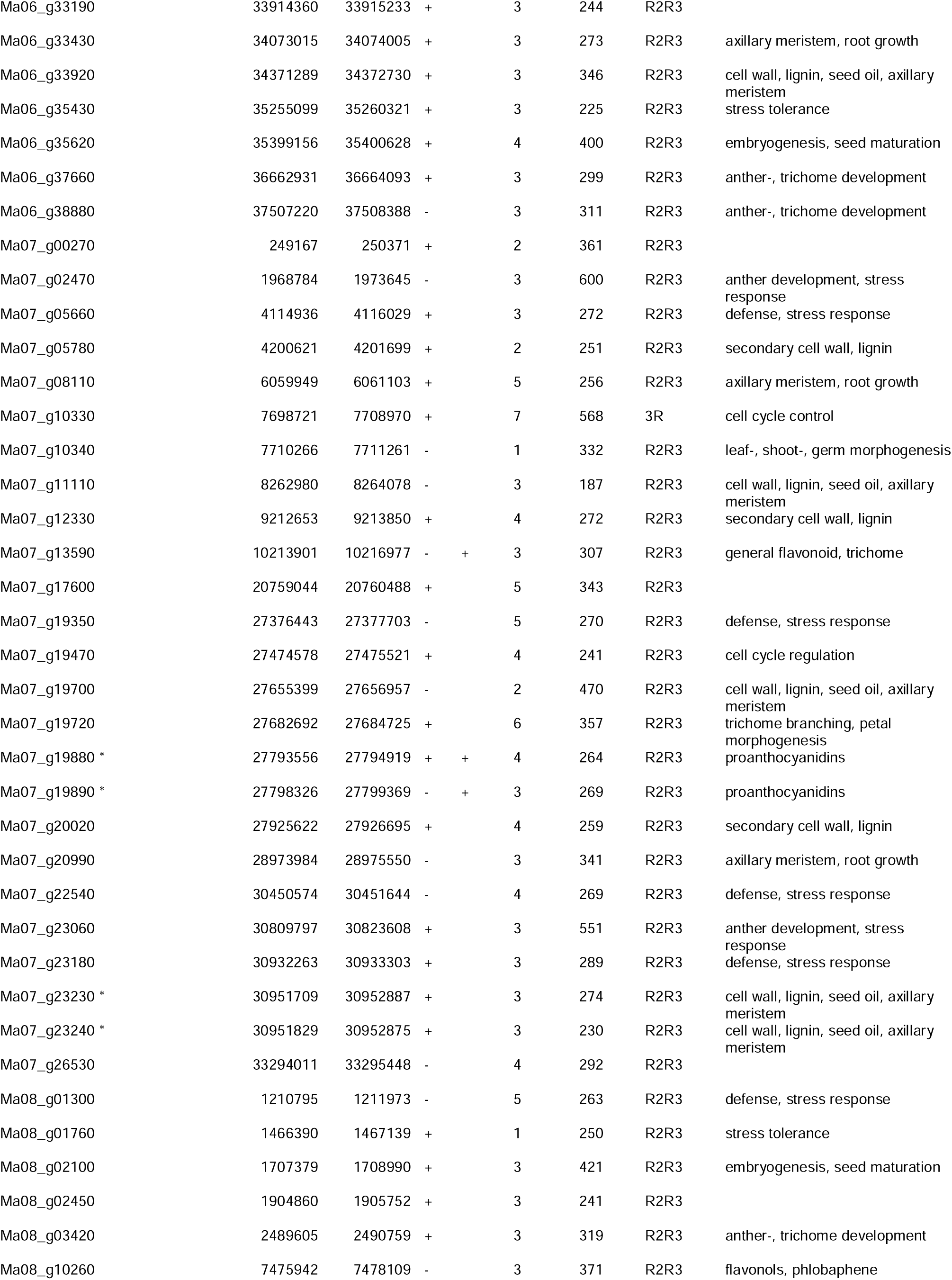
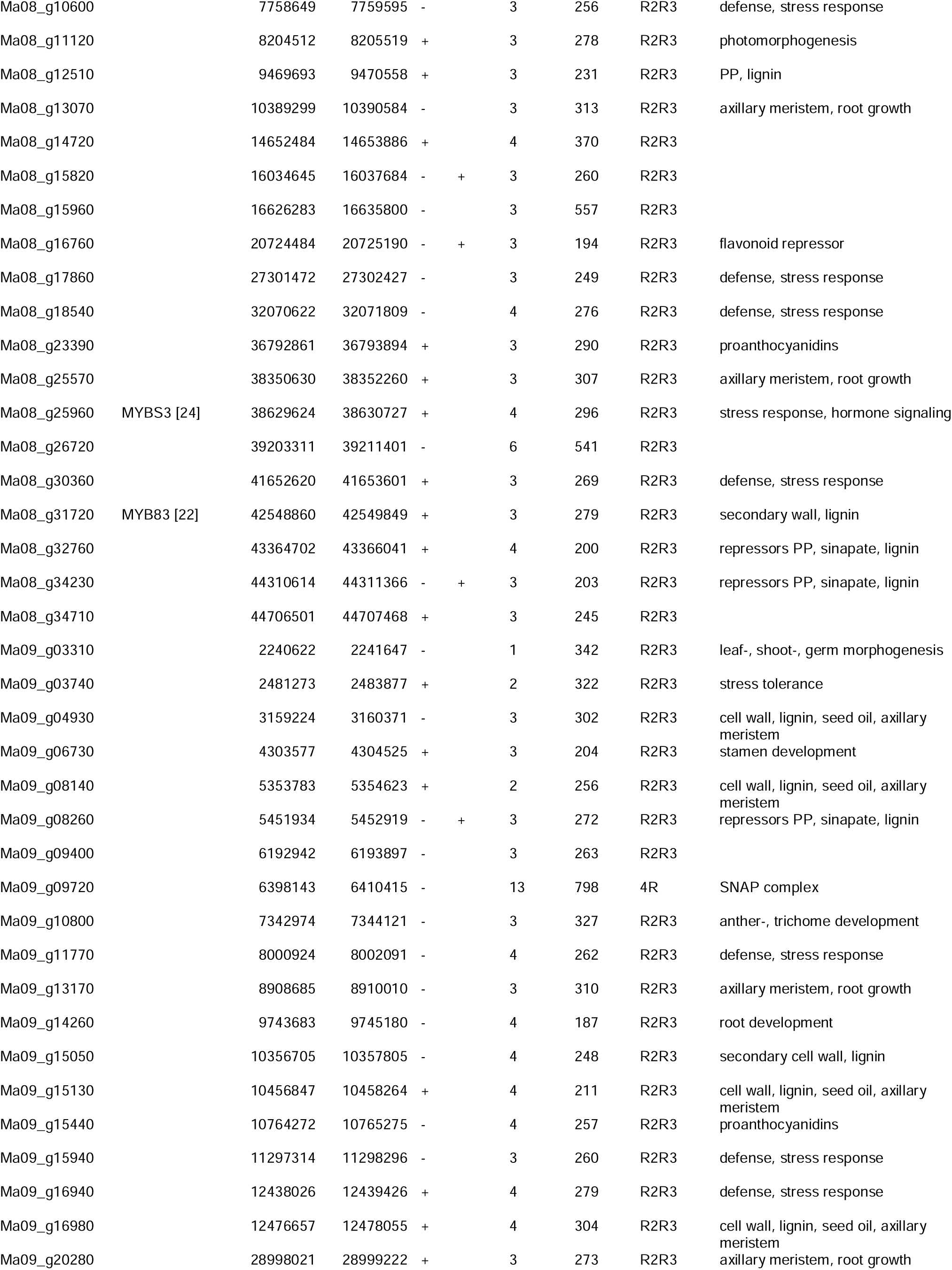
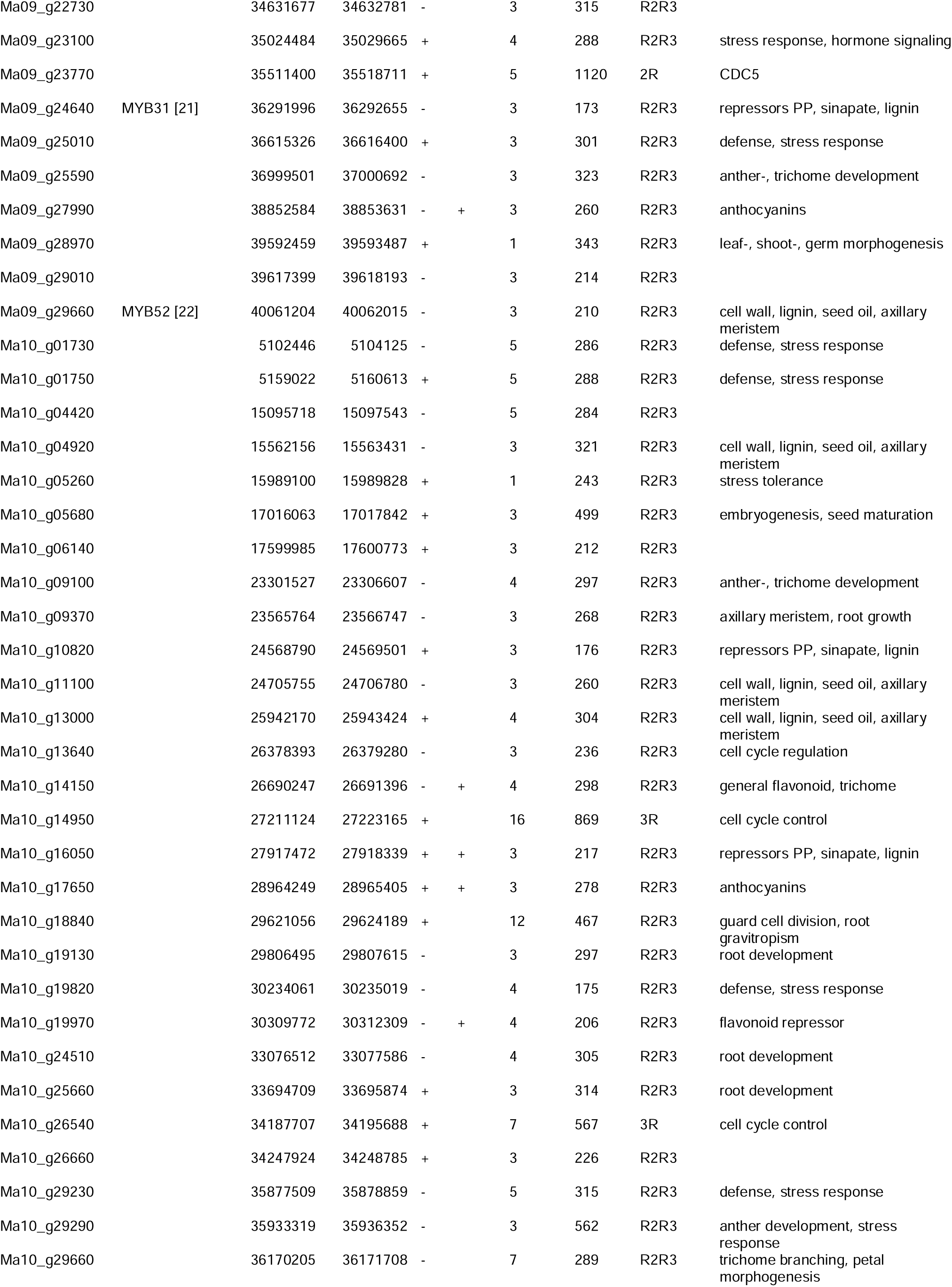
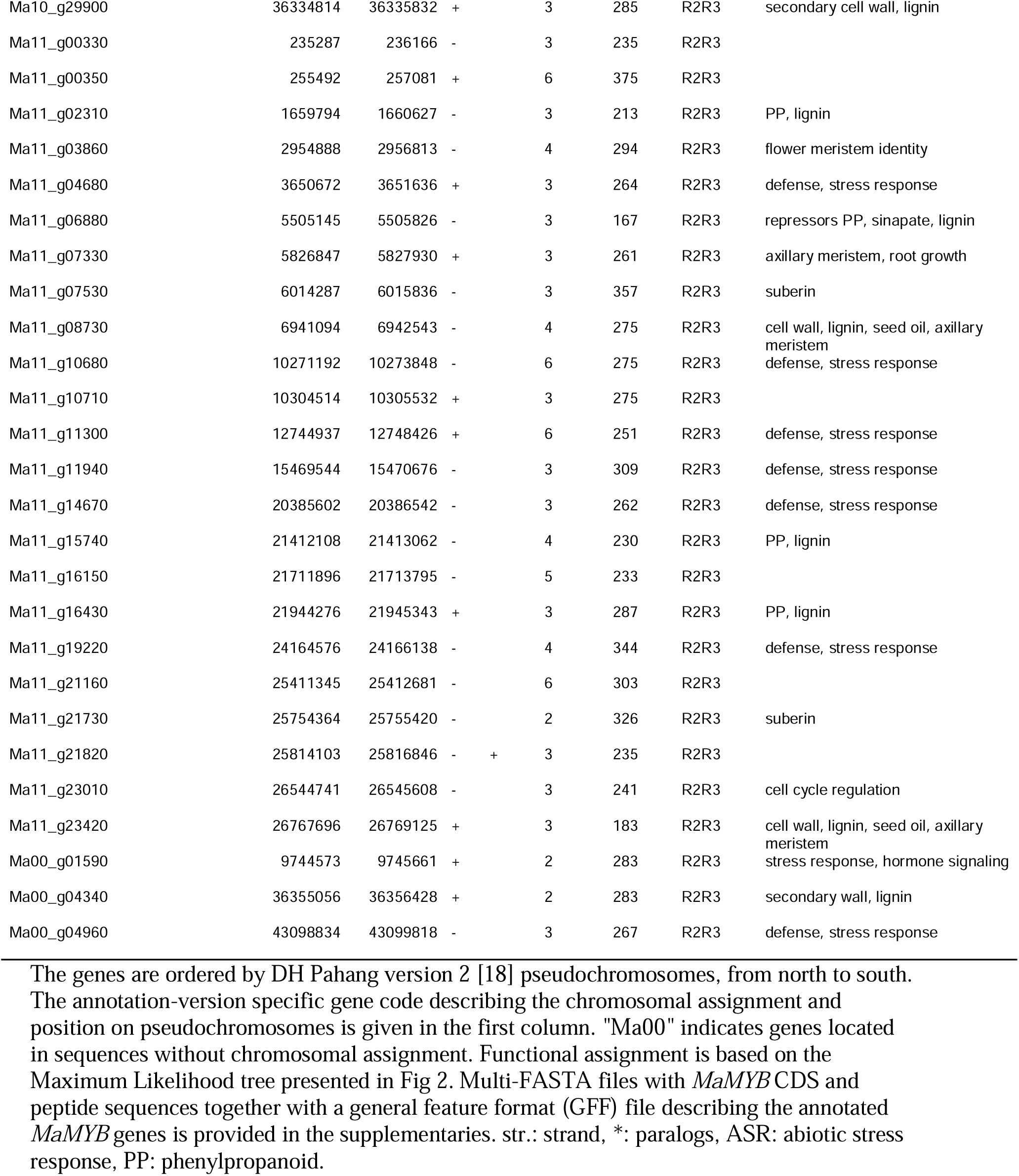
List of annotated *MYB* genes in the *Musa acuminata* (DH Pahang) genome sequence.

**Fig 1.**
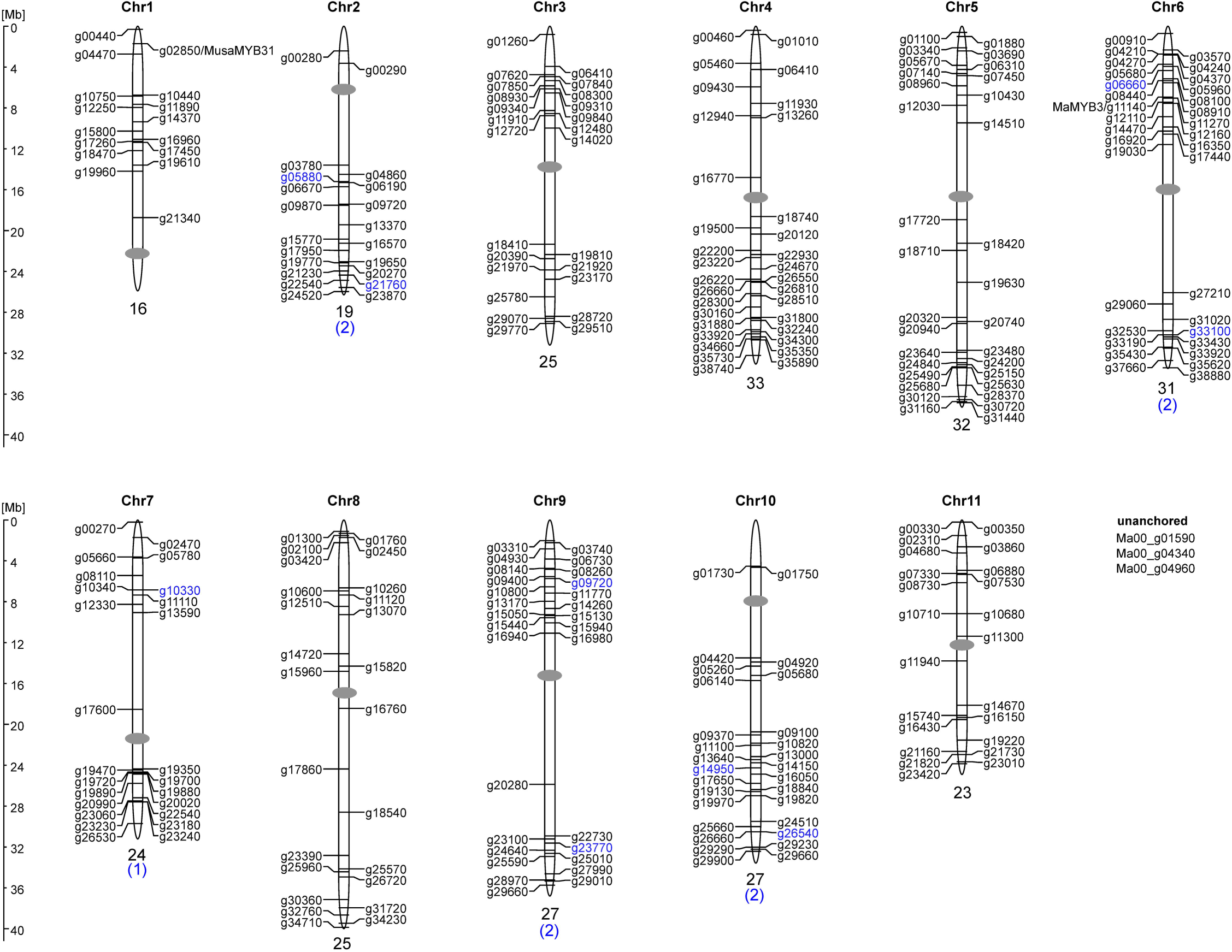
Distribution of *MaMYB* genes on the eleven *M. acuminata* chromosomes. Chromosomes are drawn to scale. The positions of centromeres (grey ovals) are roughly estimated from repeat distribution data. The chromosomal positions of the *MaMYB* genes (given in DH Pahang version 2 annotation ID) are indicated. *R2R3-MYB* genes are given in black letters, *MYB* genes with more than two MYB repeats are given in blue letters. The number of *MYB* genes on each chromosome is given below the respective chromosome.

**Fig 2.**
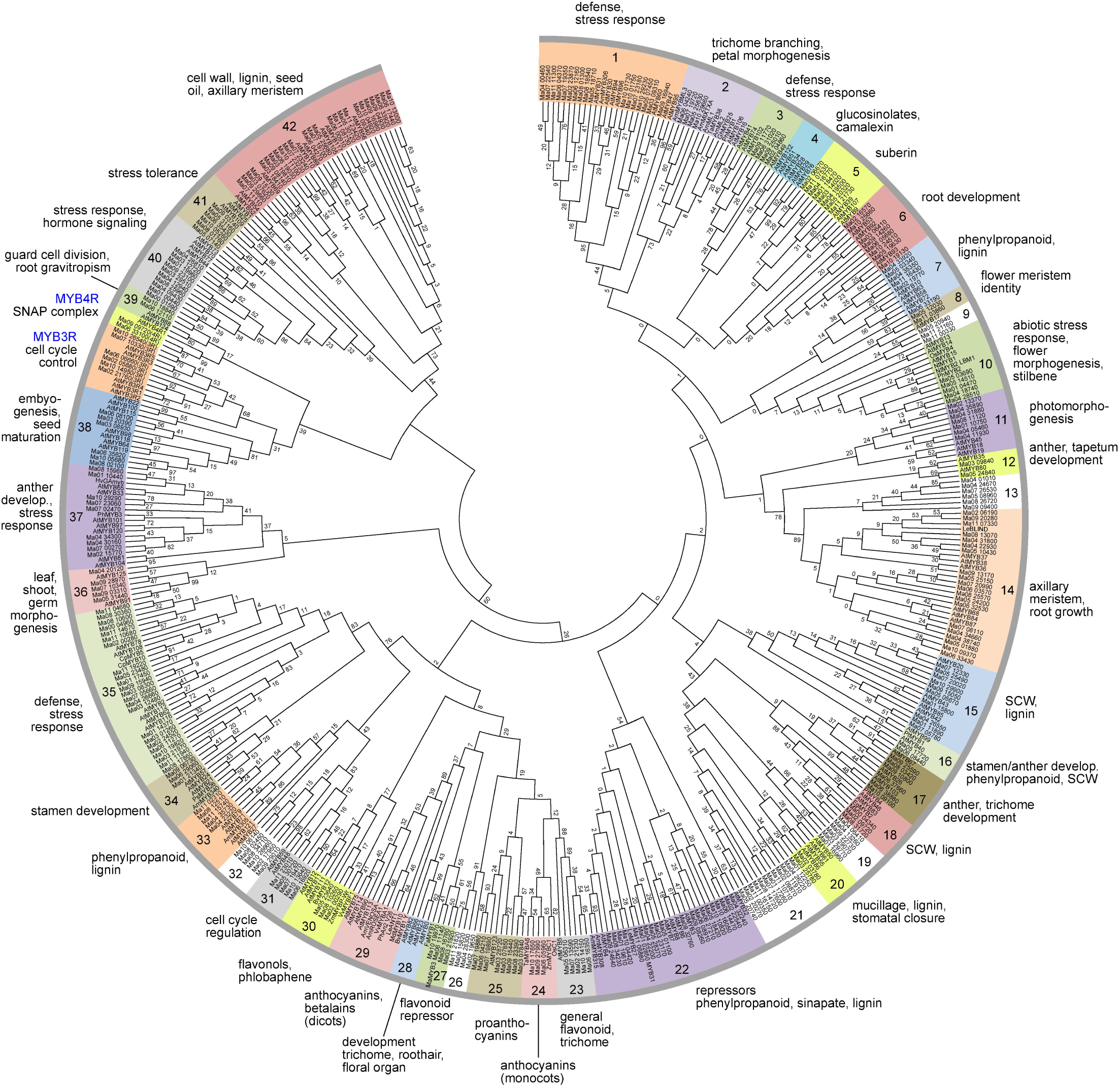
Phylogenetic Maximum Likelihood (ML) tree. ML consensus tree inferred from 1000 bootstraps with 468 MYB domain amino acid sequences of MYB proteins from *Musa accuminata* (Ma), *Arabidopsis thaliana* (At) and landmark MYBs from other plant species built with MEGA7. Clades are labeled with different colors and functional annotations are given. The numbers at the branches give bootstrap support values from 1000 replications. SCW, secondary cell wall.

The number of R2R3-MYB genes is one of the highest among the species that have been studied to date, ranging from 45 in *Ginkgo biloba* [10] over 157 in *Zea mays* [25] and 249 in *Brassica napus* [26] to 360 in *Gossypium hirsutum* [11]. This is probably due to three whole-genome duplications (γ 100 Myr ago and α, β 65 Myr ago) that occured during *Musa* genome evolution [3, 27]. The number of atypical multiple repeat *MYB* genes identified in *M. acuminata* is in the same range as those reported for most other plant species, up to six *MYB3R* and up to two *MYB4R* and *CDC5*-like genes.

The 285 *MaR2R3-MYB* genes identified constitute approximately 0.81 % of the 35,276 predicted protein-coding *M. acuminata* genes and 9.0 % of the 3,155 putative *M. acuminata* transcription factor genes [18]. These were subjected to further analyses. The identified *MaR2R3-MYB* genes were named following the nomenclature of the locus tags provided in the DH-Pahang version 2 genome annotation (Table 1). A keyword search in the NCBI database (http://www.ncbi.nlm.nih.gov/) revealed no evidence for *M. acuminata MYB* genes not present in Table 1.

Six publications dealing with *M. acuminata MYB* genes were identified: one study describes the elevated expression of nine *MaMYB* genes in transgenic banana plants overexpressing the NAC domain transcription factor MusaVND1 (vascular related NAC domain) indicating a role of these MaMYBs in the regulation of secondary wall deposition [22]. MYBS3 *(Ma08_25960)* was found to be differentially expressed between cold-sensitive and cold-tolerant bananas [24]. Another publication described the *M. acuminata R2R3-MYB* gene *Ma05_03690* being upregulated in the early response to the endoparasitic root-knot nematode *Meloidogyne incognita* in roots [28]. MusaMYB31 *(Ma01_02850)* was identified as a negative regulator of lignin biosynthesis and the general phenylpropanoid biosynthesis pathway [21]. MaMYB3 *(Ma06_11140)* was found to repress starch degradation in fruit ripening [23] and MaMYB4 *(Ma01_19610)* was recently described to control fatty acid desaturation [29].

On the basis of the DH-Pahang version 2 annotation, 291 of the 294 *MaMYB* genes could be assigned to the eleven chromosomes. The chromosomal distribution of *MaMYB* genes on the pseudochromosomes is shown in Fig 1 and revealed that *M. acuminata MYB* genes are distributed across all chromosomes.

Gene structure analysis revealed that most *MaR2R3-MYB* genes (155 of 285; 54.4 %) follow the previously reported rule of having two introns and three exons, and display the highly conserved splicing arrangement that has also been reported for other plant species [25, 30, 31]. A total of 23 (8.1 %) *MaR2R3-MYB* genes have two exons and seven (2.5 %) were one exon genes. 67 (23.5 %) *MaR2R3-MYB* genes have four exons, 21 (7.4 %) five exons, eight (2.8 %) six exons and two of each with seven and twelve exons, respectively. The complex exon-intron structure of Ma06_12110 and Ma10_18840 is conserved in their *A. thaliana* orthologs AtMYB88 and AtMYB124/FOUR LIPS (FLP) containing ten and eleven exons, respectively. This supports their close evolutionary relationship, but also suggests the conservation of this intron pattern in evolution since the monocot-dicot split 140-150 Myr ago [32].

### Phylogenetic analysis of the *M. acuminata* MYB family

With the aim to explore the putative function of the predicted *M. acuminata* MYBs, we assigned them to plant MYB proteins with known function. For this, we chose primarily data from *A. thaliana*, which is the source of most functional MYB characterisations. From comparable studies, MYB function appears conserved across MYB clades, suggesting that closely related MYBs recognise similar/same target genes and possess cooperative, overlapping or redundant functions.

To unravel the relationships, we constructed a phylogenetic tree with 468 MYB domain amino acid sequences of MYB proteins. We used 293 MaMYBs (omitting the CDC5-like MaMYB), the complete *A. thaliana* MYB family (132 members, including 126 R2R3-MYB, five MYB3R and one MYB4R) and 43 functionally well characterised landmark R2R3-MYBs from other plant species. The phylogenetic tree topology allowed us to classify the analysed MYBs into one MYB3R clade, one MYB4R clade and 42 R2R3-MYB protein clades (Fig 2).

Most R2R3-MYB clades (31 of 42) include variable numbers of MYB proteins from Arabidopsis and banana, indicating that the appearance of most *MYB* genes in these two species predates the monocot-dicot split as observed in other studies [33]. Several of these clades also contain landmark MYBs from other plant species. The two clades 4 and 28 only contain Arabidopsis R2R3-MYB members, while seven clades (9, 13, 19, 21, 26, 29 and 32, displayed with white background in Fig 2) only contain banana R2R3-MYB members. Additionally, two clades, 24 with monocot anthocyanin biosynthesis regulators and 27 with the strawberry landmark flavonoid biosynthesis repressor FaMYB1 [34], do not contain any *A. thaliana* MYB (Fig 2).

The lineage specificity of some MYB clades could indicate that these clades may have been lost or gained in a single order or species during plant evolution, as indicated by other studies [35-37]. For example, clade 28 lacked *M. acuminata* orthologs, but includes the *A. thaliana* R2R3-MYBs AtMYB0/GLABRA1 and AtMYB66/WEREWOLF, which have been identified as being involved in the formation of trichomes and root hairs from epidermal cells [38, 39]. Similar observations have been made in maize (monocot) and sugarbeet (eudicot, caryophyllales) [30, 31], both not containing clade 28 orthologs, while grape (eudicot, rosid) and poplar (eudicot, rosid) do [12, 40]. It has been hypothesized that GLABRA1-like *MYB* genes have been acquired in rosids after the rosid-asterids split [41]. The absence of MaMYBs in clade 28 is consistent with this hypothesis, since monocots branched off before the separation of asterids and rosids in eudicots. Clade 4 also lacks banana R2R3-MYBs.This clade contains the glucosinolate biosynthesis regulators AtMYB28, AtMYB34 and AtMYB51 [42, 43]. The absence of MaMYBs in this clade is concordant with the fact that glucosinolates are only present in the Brassicaceae family. This clade is thought to have originated from a duplication event before the divergence of Arabidopsis from Brassica [44].

The seven clades containing only MaMYBs were manually inspected by applying BLAST searches at the NCBI protein database in order to identify high homology to functionally characterized landmark plant R2R3-MYBs. In no case could landmark MYB be identified. Consequently, these clades could be described as a lineage-specific expansion in *M. acuminata*, reflecting a species-, genus- or order-specific evolutionary change. These MaMYB proteins may have specialised functions that were acquired or expanded in *M. acuminata* during genome evolution. Further research will be needed to decipher the biological roles of these *MaMYB* genes.

R2R3-MYBs may interact with basic helix-turn-helix (bHLH)-type transcription factors, together with WD-repeat (WDR) proteins, forming a trimeric MBW complex. These R2R3-MYBs are defined by a bHLH-binding consensus motif [D/E]Lx2[R/K]x3Lx6Lx3R [45] found in all bHLH-interacting R2R3-MYBs. A search in the MaMYB proteins for the mentioned bHLH-interaction motif identified 30 MaMYBs containing this motif (Table 1) and thus putatively interacting with bHLH proteins. 26 of these 30 MaMYBs were all functionally assigned to clades containing (potentially) known bHLH-interacting R2R3-MYBs: nine in clade 22 (repressors phenylpropanoid, sinapate-, lignin regulators), six in clade 25 (proanthocyanidin regulators), four in clade 23 (general flavonoid, trichome regulators), four in clade 27 (flavonoid repressors) and three in clade 24 (monocot anthocyanin regulators). Four MaMYBs, Ma02_19650, Ma04_28300, Ma08_15820 and Ma11_21820 were all found in the *M. acuminata*-specific clade 26. These MaMYBs may interact with bHLH-type transcription factors. The close relation to flavonoid biosynthesis regulators suggests that they may act similarly to R2R3-MYB proanthocyanidin regulators. Further research is needed to determine if they regulate a *Musa*-specific biosynthesis pathway.

### Expression profiles for *M. acuminata MYB* genes in different organs and developmental stages

Since it is not unusual for large transcription factor families in higher organisms to have redundant functions, a particular transcription factor needs to be studied and characterised in the context of the whole family. In this regard, the gene expression pattern can provide a hint to gene function. In this study we used publicly-available RNA-seq reads from the Sequence Read Archive (S2 Table) to analyse the expression of the 293 *MaMYB* genes in different organs and developmental stages: embryonic cell suspension, seedling, root and young, adult and old leaf, pulp stages S1-S4 and peel stages S1-S4. Filtered RNA-seq reads were aligned to the genome reference sequence and the number of mapped reads per annotated transcript were quantified and compared across the analysed samples to calculate normalised RNA-seq read values which are given in Table 2.

**Table 2.** The expression profiles of *MaR2R3-MYB* genes in different organs and developmental stages based on RNA-seq data.

| gene | embryonic | seedling | root | leaf | young_leaf | adult_leaf | old_leaf | pulp | pulpS1 | pulpS2 | pulpS3 | pulpS4 | peel | peelS1 | peelS2 | peelS3 | peelS4 |
| --- | --- | --- | --- | --- | --- | --- | --- | --- | --- | --- | --- | --- | --- | --- | --- | --- | --- |
| Ma00_g01590 | 50 | 35 | 6 | 151 | 77 | 175 | 348 | 6 | 5 | 5 | 8 | 5 | 8 | 2 | 19 | 10 | 2 |
| Ma00_g04340 | 0 | 0 | 0 | 0 | 0 | 0 | 0 | 0 | 1 | 0 | 0 | 0 | 0 | 0 | 0 | 0 | 0 |
| Ma00_g04960 | 0 | 0 | 0 | 0 | 0 | 0 | 0 | 0 | 0 | 0 | 0 | 0 | 0 | 0 | 0 | 0 | 0 |
| Ma01_g00440 | 181 | 49 | 38 | 129 | 53 | 175 | 276 | 32 | 31 | 58 | 34 | 7 | 30 | 9 | 63 | 35 | 11 |
| Ma01_g02850 | 9 | 2 | 14 | 27 | 21 | 31 | 38 | 20 | 24 | 23 | 23 | 10 | 11 | 7 | 9 | 17 | 12 |
| Ma01_g04470 | 1 | 0 | 12 | 4 | 1 | 6 | 2 | 9 | 13 | 9 | 7 | 8 | 6 | 6 | 3 | 7 | 7 |
| Ma01_g10440 | 3 | 16 | 4 | 5 | 7 | 5 | 5 | 4 | 5 | 5 | 3 | 3 | 3 | 3 | 2 | 3 | 4 |
| Ma01_g10750 | 0 | 0 | 2 | 2 | 0 | 0 | 4 | 1 | 1 | 1 | 1 | 2 | 4 | 5 | 4 | 4 | 5 |
| Ma01_g11890 | 0 | 0 | 3 | 3 | 10 | 1 | 0 | 0 | 0 | 0 | 0 | 0 | 0 | 0 | 0 | 0 | 0 |
| Ma01_g12250 | 1 | 0 | 4 | 5 | 17 | 2 | 0 | 3 | 4 | 3 | 2 | 2 | 4 | 4 | 4 | 3 | 3 |
| Ma01_g14370 | 0 | 17 | 11 | 0 | 0 | 0 | 0 | 5 | 7 | 7 | 8 | 0 | 8 | 3 | 7 | 18 | 4 |
| Ma01_g15800 | 1 | 0 | 4 | 1 | 0 | 0 | 0 | 4 | 2 | 5 | 7 | 3 | 6 | 8 | 5 | 2 | 9 |
| Ma01_g16960 | 0 | 0 | 1 | 0 | 1 | 0 | 0 | 0 | 0 | 0 | 0 | 0 | 0 | 0 | 0 | 0 | 0 |
| Ma01_g17260 | 1 | 0 | 11 | 1 | 0 | 0 | 0 | 3 | 3 | 3 | 2 | 3 | 4 | 5 | 3 | 4 | 5 |
| Ma01_g17450 | 0 | 0 | 6 | 2 | 0 | 1 | 6 | 5 | 13 | 2 | 2 | 1 | 1 | 1 | 2 | 2 | 0 |
| Ma01_g18470 | 0 | 0 | 0 | 0 | 0 | 0 | 0 | 0 | 1 | 0 | 0 | 0 | 1 | 2 | 1 | 0 | 0 |
| Ma01_g19610 | 13 | 0 | 6 | 39 | 56 | 40 | 57 | 29 | 78 | 22 | 15 | 2 | 12 | 7 | 36 | 2 | 5 |
| Ma01_g19960 | 35 | 1 | 6 | 97 | 94 | 148 | 145 | 5 | 7 | 6 | 5 | 4 | 19 | 5 | 58 | 10 | 4 |
| Ma01_g21340 | 0 | 0 | 0 | 0 | 0 | 0 | 0 | 2 | 8 | 0 | 0 | 0 | 0 | 0 | 0 | 0 | 0 |
| Ma02_g00280 | 1 | 1 | 21 | 4 | 1 | 1 | 5 | 6 | 8 | 10 | 4 | 2 | 5 | 5 | 4 | 5 | 7 |
| Ma02_g00290 | 0 | 0 | 3 | 9 | 5 | 26 | 1 | 8 | 25 | 0 | 6 | 2 | 3 | 2 | 3 | 2 | 5 |
| Ma02_g03780 | 0 | 0 | 29 | 18 | 26 | 0 | 0 | 13 | 14 | 9 | 18 | 12 | 30 | 39 | 31 | 17 | 33 |
| Ma02_g04860 | 0 | 0 | 1 | 0 | 0 | 0 | 0 | 0 | 0 | 1 | 0 | 0 | 0 | 0 | 0 | 0 | 0 |
| Ma02_g05880 | 4 | 2 | 2 | 3 | 2 | 5 | 3 | 2 | 4 | 2 | 2 | 1 | 2 | 1 | 1 | 1 | 3 |
| Ma02_g06190 | 0 | 0 | 19 | 8 | 0 | 0 | 0 | 16 | 30 | 12 | 13 | 11 | 28 | 42 | 18 | 18 | 32 |
| Ma02_g06670 | 1 | 1 | 4 | 1 | 0 | 0 | 0 | 4 | 3 | 7 | 4 | 4 | 5 | 6 | 2 | 6 | 5 |
| Ma02_g09720 | 0 | 0 | 0 | 0 | 0 | 0 | 0 | 0 | 0 | 0 | 0 | 0 | 0 | 1 | 0 | 0 | 1 |
| Ma02_g09870 | 29 | 1 | 23 | 37 | 99 | 33 | 11 | 19 | 15 | 28 | 27 | 6 | 11 | 11 | 10 | 9 | 13 |
| Ma02_g13370 | 0 | 0 | 1 | 0 | 0 | 0 | 0 | 0 | 0 | 0 | 0 | 0 | 0 | 0 | 0 | 0 | 0 |
| Ma02_g15770 | 6 | 0 | 1 | 0 | 0 | 1 | 0 | 1 | 2 | 0 | 1 | 0 | 1 | 1 | 1 | 0 | 1 |
| Ma02_g16570 | 0 | 0 | 0 | 0 | 1 | 0 | 1 | 0 | 0 | 0 | 0 | 0 | 0 | 0 | 0 | 0 | 0 |
| Ma02_g17950 | 8 | 0 | 46 | 14 | 4 | 6 | 13 | 33 | 53 | 35 | 16 | 26 | 19 | 19 | 14 | 20 | 22 |
| Ma02_g19650 | 0 | 0 | 3 | 3 | 5 | 1 | 0 | 7 | 10 | 5 | 3 | 11 | 9 | 15 | 8 | 6 | 5 |
| Ma02_g19770 | 0 | 0 | 0 | 0 | 0 | 0 | 0 | 0 | 0 | 0 | 0 | 0 | 0 | 0 | 0 | 0 | 0 |
| Ma02_g20270 | 3 | 0 | 3 | 16 | 57 | 4 | 0 | 1 | 1 | 2 | 2 | 1 | 4 | 3 | 4 | 3 | 7 |
| Ma02_g21230 | 2 | 2 | 1 | 5 | 14 | 3 | 1 | 3 | 8 | 3 | 0 | 0 | 1 | 0 | 3 | 2 | 0 |
| Ma02_g21760 | 0 | 3 | 7 | 2 | 2 | 0 | 0 | 8 | 11 | 7 | 8 | 7 | 8 | 11 | 5 | 5 | 12 |
| Ma02_g22540 | 1 | 0 | 7 | 4 | 2 | 6 | 7 | 1 | 1 | 1 | 2 | 0 | 1 | 1 | 1 | 1 | 1 |
| Ma02_g23870 | 3 | 0 | 0 | 7 | 4 | 20 | 3 | 2 | 8 | 0 | 0 | 0 | 0 | 0 | 0 | 0 | 0 |
| Ma02_g24520 | 0 | 0 | 0 | 0 | 0 | 0 | 0 | 0 | 1 | 0 | 0 | 0 | 0 | 0 | 0 | 0 | 0 |
| Ma03_g01260 | 11 | 0 | 6 | 16 | 0 | 0 | 61 | 9 | 31 | 3 | 1 | 2 | 2 | 1 | 3 | 4 | 1 |
| Ma03_g06410 | 0 | 0 | 11 | 1 | 0 | 0 | 0 | 1 | 0 | 3 | 0 | 1 | 1 | 2 | 0 | 2 | 1 |
| Ma03_g07620 | 0 | 0 | 3 | 0 | 0 | 0 | 0 | 0 | 0 | 0 | 0 | 0 | 0 | 0 | 0 | 0 | 0 |
| Ma03_g07840* | 0 | 0 | 2 | 1 | 2 | 0 | 0 | 10 | 24 | 9 | 3 | 5 | 2 | 1 | 3 | 1 | 2 |
| Ma03_g07850* | 0 | 0 | 1 | 1 | 1 | 0 | 0 | 1 | 2 | 1 | 0 | 1 | 2 | 2 | 2 | 3 | 1 |
| Ma03_g08300 | 0 | 1 | 3 | 0 | 0 | 0 | 1 | 1 | 5 | 0 | 0 | 0 | 1 | 0 | 0 | 6 | 0 |
| Ma03_g08930 | 0 | 0 | 0 | 0 | 0 | 0 | 0 | 0 | 0 | 0 | 0 | 0 | 0 | 0 | 0 | 0 | 0 |
| Ma03_g09310 | 0 | 0 | 0 | 12 | 42 | 7 | 1 | 0 | 0 | 0 | 0 | 0 | 0 | 0 | 0 | 0 | 0 |
| Ma03_g09340 | 2 | 0 | 7 | 1 | 1 | 0 | 1 | 5 | 8 | 4 | 5 | 2 | 3 | 4 | 2 | 3 | 5 |
| Ma03_g09840 | 0 | 0 | 0 | 0 | 0 | 0 | 0 | 0 | 0 | 0 | 0 | 0 | 0 | 0 | 0 | 0 | 0 |
| Ma03_g11910 | 5 | 8 | 25 | 35 | 76 | 25 | 6 | 12 | 10 | 13 | 12 | 13 | 24 | 26 | 30 | 18 | 22 |
| Ma03_g12480 | 0 | 0 | 4 | 0 | 0 | 0 | 0 | 1 | 1 | 1 | 0 | 0 | 0 | 0 | 0 | 1 | 0 |
| Ma03_g12720 | 16 | 10 | 4 | 45 | 59 | 49 | 70 | 2 | 2 | 2 | 2 | 3 | 18 | 2 | 32 | 35 | 3 |
| Ma03_g14020 | 0 | 0 | 1 | 0 | 0 | 0 | 0 | 2 | 1 | 4 | 1 | 3 | 2 | 3 | 1 | 3 | 2 |
| Ma03_g18410 | 0 | 0 | 0 | 1 | 2 | 0 | 0 | 4 | 15 | 0 | 0 | 0 | 0 | 0 | 0 | 0 | 0 |
| Ma03_g19810 | 0 | 0 | 2 | 0 | 0 | 0 | 0 | 0 | 0 | 0 | 0 | 0 | 0 | 0 | 0 | 0 | 0 |
| Ma03_g20390 | 5 | 1 | 1 | 2 | 2 | 0 | 4 | 2 | 1 | 3 | 2 | 2 | 0 | 1 | 0 | 0 | 1 |
| Ma03_g21920 | 5 | 0 | 2 | 17 | 13 | 32 | 22 | 4 | 12 | 2 | 2 | 0 | 2 | 0 | 6 | 3 | 0 |
| Ma03_g21970 | 0 | 0 | 1 | 0 | 0 | 0 | 0 | 0 | 0 | 0 | 0 | 0 | 0 | 0 | 0 | 0 | 0 |
| Ma03_g23170 | 0 | 0 | 3 | 4 | 8 | 2 | 0 | 2 | 2 | 3 | 1 | 2 | 4 | 6 | 2 | 3 | 4 |
| Ma03_g25780 | 3 | 2 | 60 | 18 | 14 | 3 | 0 | 37 | 28 | 81 | 18 | 22 | 40 | 57 | 29 | 26 | 48 |
| Ma03_g28720 | 3 | 3 | 1 | 2 | 0 | 1 | 5 | 1 | 3 | 1 | 0 | 1 | 0 | 0 | 0 | 0 | 0 |
| Ma03_g29070 | 0 | 0 | 1 | 2 | 4 | 2 | 0 | 1 | 1 | 0 | 3 | 0 | 2 | 1 | 2 | 1 | 2 |
| Ma03_g29510 | 14 | 4 | 5 | 44 | 49 | 65 | 55 | 9 | 25 | 5 | 5 | 2 | 6 | 3 | 7 | 11 | 3 |
| Ma03_g29770 | 1 | 0 | 3 | 1 | 1 | 0 | 0 | 2 | 3 | 4 | 0 | 2 | 1 | 2 | 1 | 1 | 2 |
| Ma04_g00460 | 2 | 0 | 1 | 9 | 14 | 20 | 2 | 0 | 0 | 0 | 0 | 0 | 1 | 0 | 2 | 1 | 0 |
| Ma04_g01010 | 0 | 0 | 7 | 0 | 0 | 0 | 1 | 1 | 3 | 1 | 1 | 0 | 1 | 1 | 1 | 0 | 1 |
| Ma04_g05460 | 0 | 0 | 0 | 0 | 0 | 0 | 0 | 0 | 0 | 0 | 1 | 0 | 1 | 0 | 1 | 0 | 1 |
| Ma04_g06410 | 0 | 1 | 1 | 0 | 1 | 0 | 0 | 0 | 1 | 0 | 0 | 0 | 0 | 0 | 0 | 0 | 0 |
| Ma04_g09430 | 4 | 1 | 9 | 12 | 17 | 19 | 2 | 4 | 4 | 2 | 6 | 5 | 8 | 11 | 7 | 3 | 11 |
| Ma04_g11930 | 0 | 0 | 0 | 0 | 0 | 0 | 0 | 0 | 0 | 0 | 0 | 0 | 0 | 0 | 0 | 0 | 0 |
| Ma04_g12940 | 4 | 0 | 2 | 3 | 6 | 2 | 4 | 2 | 3 | 1 | 2 | 0 | 1 | 0 | 1 | 0 | 0 |
| Ma04_g13260 | 0 | 2 | 2 | 1 | 1 | 1 | 2 | 2 | 3 | 2 | 3 | 1 | 0 | 0 | 0 | 0 | 0 |
| Ma04_g16770 | 0 | 1 | 1 | 1 | 0 | 1 | 0 | 1 | 0 | 1 | 0 | 3 | 1 | 1 | 0 | 1 | 0 |
| Ma04_g18740 | 11 | 1 | 7 | 2 | 1 | 1 | 5 | 2 | 0 | 0 | 6 | 0 | 1 | 0 | 1 | 2 | 0 |
| Ma04_g19500 | 0 | 0 | 4 | 1 | 0 | 0 | 0 | 2 | 4 | 3 | 0 | 1 | 1 | 1 | 1 | 1 | 0 |
| Ma04_g20120 | 0 | 0 | 0 | 0 | 0 | 0 | 0 | 0 | 0 | 0 | 0 | 0 | 0 | 0 | 0 | 0 | 0 |
| Ma04_g22200 | 0 | 0 | 3 | 0 | 0 | 0 | 0 | 3 | 2 | 8 | 1 | 3 | 1 | 1 | 0 | 1 | 0 |
| Ma04_g22930 | 0 | 0 | 4 | 1 | 0 | 0 | 0 | 2 | 2 | 3 | 0 | 2 | 4 | 6 | 2 | 3 | 4 |
| Ma04_g23220 | 0 | 0 | 5 | 3 | 0 | 0 | 1 | 1 | 3 | 1 | 1 | 1 | 4 | 3 | 6 | 1 | 6 |
| Ma04_g24670 | 0 | 0 | 33 | 6 | 0 | 0 | 0 | 24 | 32 | 24 | 19 | 19 | 17 | 22 | 9 | 16 | 19 |
| Ma04_g26220 | 0 | 1 | 1 | 2 | 1 | 1 | 0 | 1 | 3 | 1 | 1 | 0 | 4 | 2 | 11 | 2 | 1 |
| Ma04_g26550 | 0 | 0 | 5 | 1 | 1 | 0 | 0 | 2 | 3 | 3 | 1 | 1 | 1 | 1 | 1 | 1 | 3 |
| Ma04_g26660 | 0 | 2 | 22 | 1 | 0 | 1 | 3 | 12 | 27 | 11 | 5 | 4 | 1 | 0 | 2 | 1 | 2 |
| Ma04_g26810 | 41 | 78 | 15 | 153 | 258 | 145 | 185 | 39 | 35 | 74 | 32 | 17 | 33 | 13 | 54 | 50 | 14 |
| Ma04_g28300 | 0 | 0 | 0 | 0 | 0 | 0 | 0 | 0 | 0 | 0 | 0 | 0 | 0 | 0 | 0 | 0 | 0 |
| Ma04_g28510 | 8 | 1 | 3 | 11 | 2 | 22 | 10 | 9 | 13 | 8 | 14 | 2 | 3 | 2 | 5 | 3 | 3 |
| Ma04_g30160 | 1 | 0 | 0 | 0 | 1 | 1 | 0 | 2 | 5 | 1 | 0 | 0 | 0 | 0 | 0 | 0 | 0 |
| Ma04_g31800 | 0 | 0 | 0 | 1 | 0 | 0 | 3 | 0 | 1 | 1 | 0 | 0 | 2 | 0 | 1 | 6 | 0 |
| Ma04_g31880 | 0 | 0 | 2 | 0 | 0 | 0 | 0 | 1 | 0 | 1 | 1 | 0 | 1 | 1 | 2 | 0 | 1 |
| Ma04_g32240 | 0 | 0 | 1 | 0 | 0 | 0 | 0 | 2 | 2 | 4 | 2 | 0 | 0 | 1 | 0 | 0 | 1 |
| Ma04_g33920 | 3 | 7 | 2 | 2 | 4 | 1 | 1 | 0 | 1 | 1 | 0 | 0 | 1 | 1 | 1 | 3 | 1 |
| Ma04_g34300 | 1 | 0 | 0 | 1 | 2 | 1 | 0 | 2 | 4 | 1 | 1 | 2 | 1 | 1 | 0 | 0 | 1 |
| Ma04_g34660 | 0 | 1 | 0 | 0 | 0 | 0 | 0 | 1 | 0 | 0 | 0 | 3 | 0 | 0 | 0 | 0 | 0 |
| Ma04_g35350 | 0 | 0 | 0 | 0 | 0 | 0 | 0 | 0 | 0 | 0 | 0 | 0 | 0 | 0 | 0 | 0 | 0 |
| Ma04_g35730 | 0 | 0 | 4 | 0 | 0 | 0 | 0 | 0 | 1 | 0 | 0 | 0 | 0 | 0 | 0 | 0 | 0 |
| Ma04_g35890 | 1 | 0 | 0 | 0 | 0 | 0 | 0 | 1 | 2 | 1 | 0 | 0 | 0 | 1 | 0 | 1 | 0 |
| Ma04_g38740 | 0 | 0 | 1 | 0 | 0 | 0 | 0 | 0 | 0 | 0 | 0 | 0 | 1 | 1 | 1 | 0 | 1 |
| Ma05_g01100 | 0 | 0 | 0 | 0 | 0 | 0 | 1 | 0 | 0 | 0 | 0 | 0 | 0 | 0 | 0 | 0 | 0 |
| Ma05_g01880 | 0 | 0 | 2 | 1 | 0 | 0 | 0 | 2 | 3 | 2 | 0 | 2 | 2 | 1 | 1 | 2 | 2 |
| Ma05_g03340 | 0 | 0 | 2 | 1 | 1 | 0 | 1 | 1 | 0 | 2 | 0 | 0 | 0 | 0 | 0 | 0 | 0 |
| Ma05_g03690 | 10 | 0 | 1 | 14 | 2 | 25 | 28 | 1 | 0 | 1 | 1 | 1 | 2 | 0 | 8 | 1 | 0 |
| Ma05_g05670 | 1 | 0 | 1 | 0 | 0 | 0 | 0 | 1 | 2 | 0 | 0 | 0 | 0 | 0 | 0 | 1 | 0 |
| Ma05_g06310 | 4 | 1 | 1 | 4 | 4 | 4 | 8 | 1 | 3 | 1 | 1 | 0 | 0 | 0 | 1 | 0 | 0 |
| Ma05_g07140 | 4 | 0 | 0 | 0 | 0 | 0 | 0 | 0 | 1 | 1 | 1 | 0 | 0 | 0 | 0 | 2 | 0 |
| Ma05_g07450 | 4 | 0 | 0 | 2 | 4 | 3 | 1 | 1 | 2 | 1 | 1 | 0 | 0 | 0 | 0 | 0 | 0 |
| Ma05_g08960 | 27 | 1 | 9 | 34 | 47 | 53 | 23 | 17 | 38 | 11 | 8 | 10 | 9 | 10 | 3 | 9 | 12 |
| Ma05_g10430 | 0 | 0 | 2 | 1 | 0 | 0 | 0 | 2 | 1 | 1 | 2 | 3 | 3 | 6 | 2 | 1 | 4 |
| Ma05_g12030 | 4 | 1 | 2 | 1 | 1 | 1 | 2 | 3 | 8 | 2 | 1 | 2 | 2 | 2 | 2 | 1 | 1 |
| Ma05_g14510 | 1 | 0 | 4 | 48 | 0 | 1 | 190 | 1 | 0 | 1 | 2 | 0 | 1 | 1 | 1 | 3 | 0 |
| Ma05_g17720 | 0 | 0 | 1 | 0 | 0 | 0 | 0 | 0 | 1 | 0 | 0 | 0 | 4 | 6 | 4 | 2 | 4 |
| Ma05_g18420 | 0 | 0 | 0 | 0 | 0 | 1 | 0 | 1 | 4 | 0 | 0 | 0 | 0 | 0 | 0 | 0 | 0 |
| Ma05_g18710 | 4 | 0 | 3 | 3 | 4 | 4 | 5 | 3 | 5 | 2 | 3 | 0 | 2 | 1 | 4 | 2 | 1 |
| Ma05_g19630 | 0 | 0 | 6 | 0 | 0 | 0 | 0 | 1 | 2 | 3 | 0 | 1 | 1 | 1 | 0 | 2 | 0 |
| Ma05_g20320 | 1 | 0 | 2 | 10 | 5 | 14 | 19 | 0 | 0 | 1 | 0 | 0 | 1 | 1 | 0 | 1 | 0 |
| Ma05_g20740 | 0 | 4 | 1 | 2 | 9 | 0 | 0 | 2 | 5 | 1 | 2 | 0 | 1 | 0 | 2 | 1 | 0 |
| Ma05_g20940 | 1 | 0 | 5 | 11 | 0 | 0 | 44 | 1 | 1 | 0 | 1 | 0 | 0 | 1 | 0 | 0 | 1 |
| Ma05_g23480 | 0 | 0 | 0 | 0 | 0 | 0 | 0 | 2 | 3 | 5 | 0 | 0 | 0 | 0 | 0 | 0 | 1 |
| Ma05_g23640 | 0 | 0 | 1 | 1 | 0 | 3 | 0 | 2 | 7 | 0 | 2 | 0 | 2 | 1 | 3 | 0 | 2 |
| Ma05_g24200 | 0 | 0 | 0 | 0 | 0 | 0 | 0 | 0 | 1 | 0 | 0 | 0 | 0 | 0 | 0 | 0 | 0 |
| Ma05_g24840 | 0 | 0 | 0 | 0 | 0 | 0 | 0 | 0 | 0 | 0 | 0 | 0 | 0 | 0 | 0 | 0 | 0 |
| Ma05_g25150 | 0 | 0 | 6 | 2 | 0 | 0 | 0 | 3 | 4 | 4 | 5 | 1 | 3 | 3 | 2 | 2 | 4 |
| Ma05_g25490 | 0 | 1 | 2 | 2 | 1 | 3 | 1 | 1 | 2 | 2 | 1 | 1 | 1 | 2 | 1 | 1 | 2 |
| Ma05_g25630 | 0 | 0 | 0 | 4 | 15 | 0 | 1 | 1 | 6 | 0 | 0 | 0 | 0 | 0 | 0 | 0 | 0 |
| Ma05_g25680 | 1 | 0 | 1 | 2 | 2 | 1 | 3 | 7 | 24 | 2 | 3 | 1 | 0 | 0 | 1 | 1 | 0 |
| Ma05_g28370 | 0 | 0 | 4 | 4 | 9 | 4 | 1 | 1 | 0 | 1 | 0 | 1 | 2 | 2 | 1 | 1 | 2 |
| Ma05_g30120 | 0 | 3 | 16 | 1 | 0 | 1 | 0 | 4 | 7 | 4 | 2 | 2 | 2 | 2 | 1 | 2 | 2 |
| Ma05_g30720 | 0 | 0 | 19 | 1 | 0 | 0 | 0 | 7 | 12 | 10 | 2 | 4 | 3 | 3 | 3 | 1 | 6 |
| Ma05_g31160 | 9 | 3 | 5 | 6 | 7 | 7 | 7 | 5 | 7 | 4 | 4 | 3 | 2 | 2 | 3 | 2 | 4 |
| Ma05_g31440 | 0 | 3 | 2 | 68 | 266 | 2 | 0 | 43 | 12 | 88 | 71 | 2 | 26 | 2 | 89 | 9 | 2 |
| Ma06_g00910 | 0 | 7 | 1 | 5 | 1 | 1 | 15 | 0 | 0 | 0 | 0 | 0 | 1 | 1 | 1 | 3 | 0 |
| Ma06_g03570 | 0 | 0 | 4 | 2 | 0 | 0 | 0 | 3 | 5 | 3 | 3 | 4 | 5 | 7 | 4 | 3 | 7 |
| Ma06_g04210 | 0 | 0 | 5 | 2 | 1 | 0 | 0 | 5 | 5 | 2 | 7 | 4 | 3 | 4 | 3 | 2 | 5 |
| Ma06_g04240 | 1 | 0 | 1 | 1 | 4 | 0 | 0 | 7 | 29 | 0 | 0 | 0 | 0 | 0 | 0 | 0 | 0 |
| Ma06_g04270 | 11 | 1 | 1 | 1 | 0 | 1 | 2 | 3 | 6 | 3 | 2 | 1 | 0 | 0 | 1 | 0 | 0 |
| Ma06_g04370 | 7 | 0 | 10 | 13 | 5 | 29 | 15 | 1 | 3 | 0 | 0 | 0 | 2 | 1 | 6 | 1 | 0 |
| Ma06_g05680 | 0 | 0 | 7 | 4 | 0 | 0 | 0 | 20 | 15 | 34 | 15 | 16 | 8 | 9 | 2 | 11 | 10 |
| Ma06_g05960 | 0 | 0 | 6 | 4 | 4 | 0 | 0 | 13 | 9 | 19 | 13 | 9 | 4 | 4 | 3 | 9 | 2 |
| Ma06_g06660 | 3 | 2 | 2 | 3 | 2 | 3 | 6 | 2 | 3 | 1 | 2 | 2 | 1 | 1 | 1 | 1 | 1 |
| Ma06_g08100 | 3 | 0 | 0 | 0 | 0 | 0 | 0 | 0 | 0 | 0 | 0 | 0 | 0 | 0 | 0 | 0 | 0 |
| Ma06_g08440 | 3 | 0 | 1 | 0 | 0 | 0 | 0 | 0 | 0 | 0 | 0 | 0 | 0 | 1 | 0 | 0 | 0 |
| Ma06_g08910 | 4 | 3 | 5 | 28 | 68 | 23 | 16 | 8 | 6 | 11 | 9 | 5 | 10 | 4 | 20 | 14 | 3 |
| Ma06_g11140 | 4 | 0 | 1 | 14 | 6 | 33 | 16 | 2 | 4 | 2 | 0 | 1 | 1 | 0 | 4 | 1 | 0 |
| Ma06_g11270 | 0 | 1 | 10 | 0 | 0 | 0 | 0 | 0 | 1 | 0 | 0 | 0 | 1 | 0 | 1 | 3 | 0 |
| Ma06_g12110 | 0 | 1 | 1 | 1 | 0 | 1 | 1 | 1 | 2 | 1 | 2 | 1 | 1 | 1 | 1 | 1 | 2 |
| Ma06_g12160 | 1 | 1 | 1 | 1 | 1 | 2 | 2 | 1 | 2 | 0 | 1 | 1 | 1 | 0 | 0 | 2 | 1 |
| Ma06_g14470 | 0 | 0 | 3 | 1 | 1 | 1 | 2 | 2 | 1 | 2 | 2 | 2 | 1 | 0 | 1 | 3 | 0 |
| Ma06_g16350 | 0 | 1 | 0 | 3 | 4 | 2 | 5 | 1 | 2 | 1 | 1 | 2 | 1 | 0 | 0 | 1 | 0 |
| Ma06_g16920 | 3 | 1 | 40 | 38 | 30 | 13 | 13 | 60 | 108 | 51 | 34 | 46 | 54 | 72 | 36 | 45 | 61 |
| Ma06_g17440 | 0 | 0 | 0 | 0 | 0 | 0 | 0 | 1 | 1 | 1 | 0 | 1 | 1 | 0 | 1 | 1 | 2 |
| Ma06_g19030 | 0 | 0 | 2 | 0 | 0 | 0 | 1 | 3 | 8 | 1 | 1 | 1 | 1 | 1 | 0 | 1 | 0 |
| Ma06_g27210 | 2 | 2 | 8 | 6 | 2 | 6 | 16 | 3 | 4 | 5 | 0 | 2 | 2 | 2 | 1 | 2 | 2 |
| Ma06_g29060 | 9 | 2 | 0 | 0 | 0 | 0 | 1 | 2 | 7 | 1 | 0 | 0 | 0 | 0 | 0 | 0 | 0 |
| Ma06_g31020 | 7 | 3 | 7 | 6 | 3 | 9 | 8 | 1 | 2 | 2 | 2 | 0 | 2 | 1 | 7 | 1 | 1 |
| Ma06_g32530 | 7 | 0 | 0 | 2 | 1 | 4 | 2 | 0 | 0 | 0 | 0 | 0 | 0 | 0 | 0 | 0 | 1 |
| Ma06_g33100 | 4 | 6 | 3 | 3 | 2 | 3 | 3 | 4 | 5 | 3 | 2 | 6 | 3 | 4 | 2 | 2 | 5 |
| Ma06_g33190 | 0 | 0 | 0 | 1 | 1 | 1 | 1 | 0 | 0 | 0 | 0 | 0 | 0 | 0 | 0 | 0 | 0 |
| Ma06_g33430 | 0 | 1 | 11 | 2 | 1 | 1 | 0 | 8 | 20 | 7 | 2 | 4 | 3 | 5 | 3 | 4 | 3 |
| Ma06_g33920 | 0 | 7 | 10 | 4 | 3 | 0 | 1 | 5 | 5 | 4 | 3 | 6 | 10 | 13 | 8 | 9 | 9 |
| Ma06_g35430 | 6 | 15 | 28 | 31 | 49 | 14 | 31 | 20 | 37 | 22 | 13 | 8 | 33 | 24 | 63 | 25 | 17 |
| Ma06_g35620 | 0 | 0 | 0 | 0 | 0 | 0 | 0 | 0 | 0 | 0 | 0 | 0 | 0 | 0 | 1 | 0 | 0 |
| Ma06_g37660 | 0 | 0 | 0 | 0 | 0 | 0 | 0 | 0 | 0 | 0 | 0 | 0 | 0 | 0 | 0 | 0 | 0 |
| Ma06_g38880 | 1 | 0 | 0 | 1 | 2 | 0 | 0 | 0 | 0 | 0 | 0 | 0 | 0 | 0 | 0 | 0 | 0 |
| Ma07_g00270 | 0 | 0 | 1 | 1 | 1 | 0 | 0 | 0 | 1 | 0 | 0 | 0 | 0 | 1 | 0 | 0 | 0 |
| Ma07_g02470 | 5 | 1 | 4 | 5 | 5 | 5 | 7 | 5 | 6 | 7 | 2 | 3 | 3 | 3 | 2 | 2 | 4 |
| Ma07_g05660 | 0 | 0 | 0 | 0 | 0 | 0 | 0 | 0 | 0 | 0 | 0 | 0 | 1 | 0 | 0 | 2 | 0 |
| Ma07_g05780 | 0 | 1 | 1 | 3 | 6 | 2 | 1 | 2 | 3 | 2 | 2 | 1 | 2 | 1 | 1 | 2 | 2 |
| Ma07_g08110 | 0 | 1 | 5 | 1 | 0 | 0 | 4 | 0 | 1 | 0 | 0 | 0 | 0 | 0 | 0 | 1 | 0 |
| Ma07_g10330 | 2 | 4 | 4 | 1 | 1 | 1 | 2 | 2 | 1 | 2 | 1 | 2 | 2 | 2 | 1 | 1 | 3 |
| Ma07_g10340 | 2 | 11 | 6 | 23 | 82 | 3 | 2 | 11 | 22 | 10 | 8 | 5 | 14 | 8 | 15 | 28 | 5 |
| Ma07_g11110 | 0 | 0 | 0 | 3 | 5 | 5 | 1 | 0 | 0 | 0 | 0 | 0 | 0 | 0 | 0 | 0 | 0 |
| Ma07_g12330 | 0 | 0 | 1 | 7 | 5 | 18 | 3 | 1 | 2 | 1 | 0 | 0 | 0 | 0 | 1 | 0 | 0 |
| Ma07_g13590 | 0 | 0 | 1 | 6 | 4 | 16 | 3 | 0 | 1 | 0 | 0 | 0 | 0 | 0 | 0 | 0 | 0 |
| Ma07_g17600 | 1 | 1 | 8 | 2 | 4 | 0 | 1 | 1 | 1 | 1 | 0 | 1 | 2 | 2 | 2 | 2 | 2 |
| Ma07_g19350 | 2 | 0 | 1 | 22 | 18 | 44 | 23 | 4 | 15 | 0 | 0 | 1 | 9 | 1 | 26 | 11 | 0 |
| Ma07_g19470 | 3 | 0 | 15 | 7 | 5 | 6 | 6 | 54 | 80 | 62 | 33 | 42 | 10 | 14 | 6 | 9 | 10 |
| Ma07_g19700 | 0 | 0 | 0 | 0 | 0 | 0 | 0 | 1 | 4 | 0 | 0 | 0 | 0 | 0 | 0 | 0 | 0 |
| Ma07_g19720 | 0 | 0 | 7 | 7 | 26 | 1 | 0 | 5 | 11 | 6 | 3 | 1 | 4 | 2 | 12 | 1 | 1 |
| Ma07_g19880* | 1 | 1 | 5 | 4 | 0 | 0 | 2 | 10 | 14 | 7 | 11 | 8 | 9 | 12 | 3 | 8 | 12 |
| Ma07_g19890* | 0 | 0 | 0 | 0 | 0 | 0 | 0 | 0 | 2 | 0 | 0 | 0 | 0 | 0 | 0 | 0 | 0 |
| Ma07_g20020 | 2 | 0 | 1 | 0 | 0 | 1 | 1 | 0 | 0 | 1 | 0 | 0 | 0 | 0 | 1 | 1 | 0 |
| Ma07_g20990 | 0 | 0 | 2 | 0 | 0 | 0 | 0 | 2 | 2 | 1 | 3 | 0 | 2 | 2 | 1 | 1 | 1 |
| Ma07_g22540 | 5 | 1 | 0 | 8 | 13 | 12 | 7 | 1 | 3 | 0 | 0 | 0 | 1 | 0 | 3 | 1 | 0 |
| Ma07_g23060 | 0 | 1 | 2 | 1 | 2 | 0 | 0 | 2 | 4 | 2 | 3 | 1 | 1 | 2 | 1 | 1 | 1 |
| Ma07_g23180 | 3 | 1 | 1 | 3 | 8 | 1 | 0 | 4 | 14 | 1 | 1 | 0 | 3 | 4 | 2 | 1 | 5 |
| Ma07_g23230* | 0 | 171 | 9 | 2 | 2 | 1 | 3 | 34 | 6 | 10 | 12 | 106 | 9 | 1 | 2 | 33 | 2 |
| Ma07_g23240* | 16 | 23 | 7 | 9 | 9 | 12 | 11 | 9 | 7 | 10 | 12 | 6 | 5 | 6 | 3 | 3 | 7 |
| Ma07_g26530 | 0 | 1 | 1 | 1 | 0 | 2 | 1 | 1 | 4 | 0 | 0 | 0 | 0 | 0 | 0 | 0 | 0 |
| Ma08_g01300 | 6 | 0 | 3 | 6 | 12 | 3 | 7 | 0 | 0 | 0 | 0 | 0 | 1 | 0 | 0 | 2 | 0 |
| Ma08_g01760 | 6 | 0 | 0 | 2 | 2 | 2 | 4 | 0 | 1 | 0 | 0 | 0 | 1 | 0 | 1 | 1 | 1 |
| Ma08_g02100 | 0 | 0 | 0 | 0 | 0 | 0 | 0 | 0 | 0 | 0 | 0 | 0 | 0 | 0 | 0 | 0 | 0 |
| Ma08_g02450 | 1 | 3 | 5 | 2 | 2 | 1 | 3 | 7 | 7 | 10 | 9 | 1 | 5 | 3 | 13 | 2 | 3 |
| Ma08_g03420 | 0 | 0 | 1 | 0 | 0 | 0 | 0 | 0 | 0 | 0 | 0 | 0 | 0 | 0 | 0 | 0 | 0 |
| Ma08_g10260 | 1 | 0 | 3 | 7 | 16 | 3 | 1 | 4 | 12 | 1 | 1 | 3 | 13 | 18 | 6 | 7 | 19 |
| Ma08_g10600 | 0 | 0 | 0 | 0 | 0 | 0 | 1 | 0 | 0 | 1 | 1 | 0 | 0 | 0 | 0 | 1 | 0 |
| Ma08_g11120 | 0 | 0 | 0 | 0 | 0 | 0 | 0 | 0 | 1 | 1 | 0 | 0 | 0 | 0 | 0 | 0 | 0 |
| Ma08_g12510 | 0 | 0 | 3 | 1 | 0 | 0 | 0 | 2 | 3 | 2 | 2 | 0 | 2 | 2 | 1 | 1 | 5 |
| Ma08_g13070 | 0 | 0 | 10 | 4 | 0 | 0 | 0 | 17 | 12 | 30 | 18 | 8 | 15 | 19 | 7 | 12 | 21 |
| Ma08_g14720 | 0 | 1 | 5 | 5 | 14 | 1 | 0 | 2 | 4 | 1 | 3 | 2 | 6 | 8 | 6 | 2 | 7 |
| Ma08_g15820 | 0 | 1 | 4 | 2 | 1 | 1 | 2 | 6 | 7 | 6 | 6 | 4 | 4 | 5 | 4 | 5 | 2 |
| Ma08_g15960 | 2 | 6 | 3 | 1 | 2 | 1 | 1 | 4 | 5 | 1 | 2 | 5 | 2 | 2 | 1 | 1 | 2 |
| Ma08_g16760 | 2 | 0 | 2 | 10 | 5 | 13 | 21 | 3 | 5 | 3 | 0 | 3 | 3 | 1 | 4 | 7 | 0 |
| Ma08_g17860 | 0 | 0 | 1 | 2 | 0 | 0 | 8 | 0 | 0 | 0 | 0 | 0 | 2 | 0 | 0 | 7 | 0 |
| Ma08_g18540 | 0 | 0 | 0 | 1 | 0 | 1 | 2 | 0 | 0 | 0 | 0 | 0 | 0 | 0 | 0 | 2 | 0 |
| Ma08_g23390 | 0 | 0 | 0 | 0 | 0 | 0 | 0 | 2 | 6 | 0 | 0 | 0 | 0 | 0 | 1 | 0 | 0 |
| Ma08_g25570 | 7 | 1 | 16 | 11 | 7 | 6 | 7 | 20 | 20 | 26 | 25 | 10 | 19 | 16 | 19 | 16 | 23 |
| Ma08_g25960 | 9 | 11 | 9 | 21 | 22 | 32 | 15 | 13 | 18 | 17 | 11 | 5 | 9 | 9 | 9 | 5 | 13 |
| Ma08_g26720 | 1 | 0 | 7 | 9 | 19 | 4 | 13 | 1 | 1 | 2 | 1 | 1 | 1 | 0 | 1 | 1 | 1 |
| Ma08_g30360 | 0 | 0 | 5 | 0 | 0 | 0 | 0 | 1 | 4 | 0 | 0 | 0 | 1 | 0 | 0 | 3 | 0 |
| Ma08_g31720 | 0 | 0 | 1 | 1 | 1 | 0 | 0 | 0 | 0 | 0 | 1 | 0 | 0 | 0 | 0 | 0 | 0 |
| Ma08_g32760 | 1 | 3 | 11 | 4 | 4 | 1 | 1 | 5 | 5 | 4 | 9 | 3 | 9 | 8 | 16 | 5 | 7 |
| Ma08_g34230 | 0 | 0 | 2 | 1 | 0 | 1 | 0 | 2 | 4 | 1 | 1 | 1 | 2 | 1 | 1 | 2 | 2 |
| Ma08_g34710 | 0 | 0 | 15 | 1 | 0 | 0 | 0 | 16 | 17 | 20 | 8 | 20 | 5 | 5 | 4 | 6 | 7 |
| Ma09_g03310 | 2 | 1 | 11 | 4 | 8 | 3 | 1 | 2 | 5 | 1 | 1 | 2 | 3 | 3 | 5 | 1 | 3 |
| Ma09_g03740 | 8 | 1 | 3 | 4 | 5 | 5 | 4 | 4 | 6 | 3 | 3 | 4 | 2 | 2 | 2 | 2 | 2 |
| Ma09_g04930 | 0 | 2 | 12 | 2 | 0 | 0 | 0 | 10 | 7 | 22 | 7 | 3 | 5 | 5 | 9 | 4 | 3 |
| Ma09_g06730 | 1 | 0 | 0 | 0 | 0 | 0 | 0 | 1 | 1 | 1 | 0 | 0 | 0 | 0 | 0 | 0 | 0 |
| Ma09_g08140 | 0 | 0 | 2 | 0 | 1 | 0 | 0 | 3 | 10 | 0 | 0 | 0 | 0 | 0 | 0 | 0 | 0 |
| Ma09_g08260 | 15 | 1 | 9 | 6 | 8 | 4 | 3 | 9 | 9 | 13 | 10 | 4 | 8 | 9 | 8 | 6 | 8 |
| Ma09_g09400 | 0 | 0 | 7 | 1 | 0 | 0 | 1 | 1 | 2 | 2 | 0 | 1 | 1 | 0 | 1 | 1 | 0 |
| Ma09_g09720 | 1 | 0 | 3 | 1 | 0 | 1 | 0 | 2 | 2 | 2 | 2 | 1 | 1 | 1 | 0 | 1 | 1 |
| Ma09_g10800 | 0 | 0 | 5 | 0 | 0 | 0 | 0 | 1 | 1 | 2 | 0 | 2 | 1 | 1 | 0 | 1 | 0 |
| Ma09_g11770 | 2 | 0 | 0 | 0 | 0 | 0 | 0 | 0 | 0 | 0 | 0 | 0 | 0 | 0 | 0 | 0 | 0 |
| Ma09_g13170 | 0 | 0 | 5 | 3 | 0 | 0 | 0 | 6 | 7 | 4 | 10 | 4 | 9 | 12 | 4 | 5 | 13 |
| Ma09_g14260 | 0 | 0 | 1 | 0 | 0 | 0 | 0 | 0 | 0 | 0 | 0 | 0 | 1 | 3 | 0 | 2 | 1 |
| Ma09_g15050 | 1 | 0 | 0 | 0 | 0 | 0 | 0 | 1 | 0 | 0 | 1 | 1 | 1 | 1 | 0 | 0 | 0 |
| Ma09_g15130 | 1 | 0 | 1 | 0 | 0 | 0 | 0 | 1 | 0 | 0 | 3 | 0 | 1 | 1 | 0 | 1 | 1 |
| Ma09_g15440 | 0 | 0 | 1 | 2 | 5 | 1 | 0 | 5 | 9 | 3 | 2 | 4 | 1 | 1 | 2 | 2 | 1 |
| Ma09_g15940 | 2 | 0 | 2 | 3 | 0 | 0 | 13 | 0 | 0 | 0 | 0 | 0 | 0 | 0 | 0 | 0 | 0 |
| Ma09_g16940 | 3 | 0 | 0 | 4 | 5 | 8 | 2 | 0 | 1 | 0 | 0 | 0 | 1 | 2 | 1 | 0 | 1 |
| Ma09_g16980 | 0 | 1 | 7 | 0 | 0 | 0 | 0 | 1 | 4 | 1 | 0 | 1 | 1 | 1 | 1 | 1 | 1 |
| Ma09_g20280 | 0 | 0 | 62 | 8 | 0 | 0 | 0 | 44 | 31 | 73 | 45 | 27 | 26 | 30 | 21 | 29 | 23 |
| Ma09_g22730 | 0 | 0 | 2 | 0 | 0 | 0 | 0 | 1 | 1 | 0 | 0 | 1 | 1 | 1 | 1 | 1 | 1 |
| Ma09_g23100 | 32 | 2 | 3 | 36 | 45 | 65 | 32 | 10 | 9 | 21 | 9 | 1 | 5 | 2 | 12 | 4 | 3 |
| Ma09_g24640 | 0 | 0 | 8 | 4 | 3 | 0 | 8 | 7 | 12 | 6 | 3 | 6 | 4 | 5 | 2 | 6 | 4 |
| Ma09_g25010 | 4 | 0 | 2 | 4 | 3 | 4 | 7 | 1 | 1 | 1 | 2 | 0 | 2 | 1 | 2 | 1 | 2 |
| Ma09_g25590 | 0 | 0 | 2 | 0 | 0 | 0 | 0 | 0 | 0 | 0 | 0 | 0 | 0 | 0 | 0 | 0 | 0 |
| Ma09_g27990 | 0 | 0 | 1 | 1 | 2 | 0 | 0 | 1 | 2 | 2 | 0 | 1 | 2 | 2 | 1 | 1 | 3 |
| Ma09_g28970 | 1 | 0 | 0 | 7 | 26 | 2 | 0 | 1 | 4 | 0 | 0 | 0 | 0 | 0 | 0 | 0 | 0 |
| Ma09_g29010 | 0 | 2 | 32 | 6 | 0 | 1 | 3 | 25 | 26 | 32 | 15 | 25 | 15 | 16 | 7 | 17 | 22 |
| Ma09_g29660 | 0 | 0 | 0 | 0 | 0 | 2 | 0 | 0 | 0 | 0 | 0 | 0 | 0 | 0 | 0 | 0 | 0 |
| Ma10_g01730 | 0 | 0 | 0 | 1 | 4 | 1 | 0 | 0 | 0 | 0 | 0 | 0 | 0 | 0 | 0 | 0 | 0 |
| Ma10_g01750 | 0 | 0 | 0 | 0 | 1 | 1 | 0 | 0 | 0 | 0 | 0 | 0 | 0 | 0 | 0 | 0 | 0 |
| Ma10_g04420 | 0 | 0 | 1 | 4 | 10 | 2 | 2 | 1 | 2 | 1 | 1 | 0 | 1 | 1 | 3 | 0 | 0 |
| Ma10_g04920 | 3 | 0 | 0 | 1 | 3 | 2 | 0 | 0 | 1 | 0 | 0 | 0 | 0 | 0 | 1 | 0 | 0 |
| Ma10_g05260 | 0 | 0 | 0 | 1 | 1 | 1 | 3 | 0 | 0 | 0 | 0 | 0 | 0 | 0 | 0 | 0 | 0 |
| Ma10_g05680 | 0 | 0 | 0 | 0 | 0 | 0 | 0 | 0 | 0 | 0 | 1 | 0 | 0 | 0 | 0 | 0 | 0 |
| Ma10_g06140 | 0 | 0 | 0 | 0 | 0 | 0 | 1 | 0 | 1 | 0 | 0 | 0 | 0 | 0 | 0 | 0 | 0 |
| Ma10_g09100 | 0 | 0 | 0 | 1 | 4 | 0 | 0 | 0 | 0 | 0 | 0 | 0 | 0 | 0 | 0 | 0 | 0 |
| Ma10_g09370 | 0 | 0 | 4 | 3 | 1 | 0 | 5 | 2 | 2 | 3 | 3 | 1 | 3 | 3 | 2 | 4 | 2 |
| Ma10_g10820 | 0 | 0 | 2 | 0 | 0 | 0 | 0 | 4 | 14 | 1 | 0 | 1 | 1 | 2 | 1 | 1 | 1 |
| Ma10_g11100 | 0 | 1 | 6 | 2 | 0 | 0 | 6 | 0 | 0 | 0 | 0 | 0 | 2 | 0 | 0 | 7 | 0 |
| Ma10_g13000 | 3 | 0 | 5 | 10 | 2 | 21 | 16 | 1 | 1 | 1 | 0 | 2 | 1 | 1 | 1 | 1 | 1 |
| Ma10_g13640 | 2 | 1 | 40 | 31 | 30 | 34 | 12 | 22 | 25 | 19 | 20 | 25 | 35 | 48 | 21 | 28 | 44 |
| Ma10_g14150 | 0 | 1 | 1 | 2 | 2 | 3 | 4 | 1 | 3 | 1 | 0 | 1 | 1 | 0 | 1 | 1 | 0 |
| Ma10_g14950 | 0 | 0 | 3 | 2 | 0 | 0 | 0 | 4 | 8 | 1 | 2 | 3 | 4 | 4 | 2 | 3 | 7 |
| Ma10_g16050 | 5 | 1 | 10 | 28 | 74 | 23 | 13 | 9 | 14 | 16 | 4 | 3 | 7 | 4 | 16 | 3 | 4 |
| Ma10_g17650 | 0 | 30 | 0 | 0 | 0 | 0 | 0 | 4 | 13 | 1 | 0 | 0 | 2 | 2 | 2 | 4 | 1 |
| Ma10_g18840 | 3 | 1 | 5 | 2 | 1 | 2 | 2 | 4 | 6 | 3 | 6 | 2 | 3 | 5 | 2 | 2 | 5 |
| Ma10_g19130 | 0 | 0 | 2 | 0 | 0 | 0 | 0 | 1 | 0 | 0 | 1 | 1 | 1 | 0 | 1 | 0 | 1 |
| Ma10_g19820 | 1 | 0 | 2 | 6 | 0 | 0 | 22 | 0 | 0 | 0 | 0 | 0 | 5 | 0 | 12 | 9 | 0 |
| Ma10_g19970 | 1 | 0 | 0 | 2 | 6 | 3 | 1 | 1 | 3 | 1 | 0 | 0 | 0 | 0 | 0 | 0 | 0 |
| Ma10_g24510 | 9 | 4 | 2 | 7 | 16 | 6 | 4 | 3 | 4 | 2 | 4 | 2 | 1 | 1 | 1 | 1 | 2 |
| Ma10_g25660 | 1 | 0 | 0 | 0 | 0 | 0 | 0 | 1 | 1 | 0 | 0 | 1 | 0 | 0 | 0 | 0 | 0 |
| Ma10_g26540 | 5 | 7 | 6 | 4 | 3 | 4 | 4 | 6 | 8 | 4 | 4 | 6 | 4 | 5 | 2 | 3 | 6 |
| Ma10_g26660 | 2 | 0 | 6 | 3 | 1 | 2 | 2 | 10 | 14 | 11 | 8 | 5 | 4 | 3 | 3 | 5 | 6 |
| Ma10_g29230 | 6 | 0 | 1 | 4 | 6 | 9 | 3 | 0 | 1 | 1 | 0 | 0 | 1 | 0 | 2 | 0 | 0 |
| Ma10_g29290 | 4 | 1 | 1 | 2 | 2 | 2 | 2 | 1 | 1 | 0 | 1 | 1 | 1 | 1 | 0 | 0 | 1 |
| Ma10_g29660 | 2 | 0 | 1 | 15 | 60 | 0 | 0 | 1 | 3 | 0 | 0 | 0 | 0 | 0 | 0 | 0 | 0 |
| Ma10_g29900 | 5 | 1 | 9 | 10 | 7 | 11 | 11 | 8 | 8 | 7 | 9 | 8 | 9 | 11 | 10 | 10 | 7 |
| Ma11_g00330 | 0 | 0 | 4 | 0 | 0 | 0 | 0 | 0 | 1 | 0 | 0 | 1 | 0 | 0 | 0 | 0 | 0 |
| Ma11_g00350 | 0 | 0 | 6 | 5 | 13 | 1 | 0 | 3 | 5 | 2 | 2 | 2 | 5 | 7 | 5 | 3 | 6 |
| Ma11_g02310 | 0 | 0 | 2 | 1 | 0 | 0 | 0 | 2 | 1 | 1 | 4 | 0 | 2 | 2 | 1 | 1 | 4 |
| Ma11_g03860 | 0 | 0 | 2 | 13 | 47 | 1 | 0 | 5 | 14 | 4 | 1 | 0 | 2 | 1 | 5 | 1 | 1 |
| Ma11_g04680 | 20 | 2 | 0 | 32 | 6 | 57 | 64 | 0 | 1 | 1 | 0 | 0 | 4 | 0 | 7 | 10 | 0 |
| Ma11_g06880 | 22 | 4 | 15 | 13 | 7 | 19 | 12 | 31 | 41 | 39 | 30 | 14 | 7 | 6 | 4 | 9 | 11 |
| Ma11_g07330 | 0 | 0 | 0 | 0 | 0 | 0 | 0 | 0 | 0 | 0 | 0 | 0 | 0 | 0 | 0 | 0 | 0 |
| Ma11_g07530 | 0 | 0 | 5 | 2 | 1 | 1 | 1 | 6 | 3 | 14 | 4 | 2 | 3 | 5 | 1 | 4 | 2 |
| Ma11_g08730 | 1 | 1 | 1 | 1 | 1 | 1 | 1 | 1 | 1 | 1 | 0 | 1 | 1 | 0 | 1 | 0 | 1 |
| Ma11_g10680 | 1 | 2 | 5 | 2 | 1 | 0 | 3 | 12 | 21 | 20 | 6 | 3 | 6 | 7 | 5 | 8 | 6 |
| Ma11_g10710 | 0 | 1 | 2 | 1 | 4 | 0 | 0 | 2 | 5 | 2 | 1 | 2 | 3 | 1 | 6 | 2 | 1 |
| Ma11_g11300 | 2 | 0 | 0 | 0 | 0 | 0 | 1 | 0 | 0 | 0 | 0 | 0 | 0 | 0 | 0 | 1 | 0 |
| Ma11_g11940 | 0 | 0 | 5 | 0 | 0 | 0 | 0 | 0 | 1 | 0 | 0 | 0 | 0 | 0 | 0 | 0 | 0 |
| Ma11_g14670 | 0 | 1 | 2 | 0 | 0 | 0 | 0 | 0 | 0 | 0 | 0 | 0 | 2 | 0 | 2 | 5 | 0 |
| Ma11_g15740 | 4 | 0 | 2 | 5 | 7 | 4 | 6 | 6 | 9 | 4 | 6 | 5 | 2 | 1 | 3 | 2 | 1 |
| Ma11_g16150 | 1 | 2 | 3 | 10 | 37 | 2 | 1 | 4 | 7 | 4 | 6 | 1 | 3 | 3 | 3 | 2 | 4 |
| Ma11_g16430 | 0 | 0 | 0 | 0 | 0 | 0 | 0 | 0 | 0 | 0 | 0 | 0 | 0 | 0 | 0 | 0 | 0 |
| Ma11_g19220 | 1 | 0 | 1 | 0 | 0 | 0 | 0 | 0 | 1 | 1 | 0 | 0 | 2 | 0 | 0 | 7 | 0 |
| Ma11_g21160 | 1 | 0 | 9 | 3 | 0 | 1 | 12 | 2 | 4 | 0 | 4 | 0 | 0 | 1 | 0 | 0 | 0 |
| Ma11_g21730 | 0 | 0 | 2 | 1 | 0 | 1 | 0 | 2 | 2 | 3 | 0 | 1 | 2 | 4 | 0 | 2 | 2 |
| Ma11_g21820 | 0 | 0 | 0 | 2 | 2 | 4 | 0 | 0 | 0 | 0 | 0 | 0 | 0 | 0 | 1 | 0 | 0 |
| Ma11_g23010 | 0 | 0 | 6 | 1 | 0 | 0 | 0 | 3 | 3 | 5 | 2 | 2 | 2 | 3 | 1 | 2 | 3 |
| Ma11_g23420 | 1 | 0 | 2 | 0 | 0 | 0 | 0 | 1 | 2 | 0 | 0 | 0 | 0 | 0 | 0 | 0 | 0 |

The intensity of green background is correlated to expression level given in FPKM. \* indicate paralogs.

Our expression analyses revealed that *M. acuminata* MYBs have diverse expression patterns in different organs. Many of the *MaMYBs* exhibited low transcript abundance levels with expression in only one or a few organs. This is consistent with other transcription factor genes, typically found to be expressed in this manner due to functional specificity and diversity. The highest number of expressed *MaMYB* genes (221; 75.4%) is observed in roots, followed by pulp (209; 71.3%), leaf (203; 69.3%), peel (196; 66.9%) and embryonic cell suspension (129; 44%). The fewest *MaMYB* genes are expressed in seedlings (98; 33.4%) in the considered dataset. 40 *MaMYB* genes (13.7%) were expressed in all samples analysed (albeit with varying expression levels), which suggested that these MaMYBs play regulatory roles at multiple developmental stages in multiple tissues. 13 *MaMYB* genes (4.4%) lacked expression information in any of the analysed samples, possibly indicating that these genes are expressed in other organs (e.g. pseudostem, flower, bract), specific cells, at specific developmental stages, under special conditions or are pseudogenes. 280 MaMYBs (95.6%) are expressed in at least one analysed sample, although the transcript abundance of some genes was very low. Some *MaMYB* genes were expressed in all analysed RNA-seq samples at similar levels (e.g. R2R3-MYBs *Ma06_33440* and *Ma11_06880*) while others show variance in transcript abundance with low (no) levels in one or several organs and high levels in others (or vice versa). For example, *Ma02_16570, Ma03_09310, Ma07_11110, Ma10_01730, Ma10_05260* and *Ma10_09100* show organ-specific expression, as their transcripts were exclusively detected in leaves, which hints to leaf-specific functionality. *Ma03_07840* and *Ma07_19470* were found to be predominantly expressed in pulp, showing expression also in other analysed organs, but not in the seedling. Overall, these results suggests that the corresponding MaMYB regulators are limited to distinct organs, tissues, cells or conditions.

Some paralogous *MaMYB* genes clustered in the genome (Tables 1 and 2) showed different expression profiles, while other clustered paralogous *MaMYB* genes did not. For example, the clustered *R2R3-MYB* genes *Ma07_19880* and *Ma07_19890* (both in the proanthocyanidin-related cluster 25): while *Ma07_19880* is expressed in pulps, peels, roots, and leaves, *Ma07_19890* is nearly not expressed in the analysed organs. The expression pattern of *Ma03_07840* and *Ma03_07850* (also both in the proanthocyanidin-related cluster 25) is, in contrast, very similar, with low expression in root, leaf, pulp and peel, but no expression in embyonic cells and roots. These results could point to functional redundancy of the genes *Ma03_07840* and *Ma03_07850*, while *Ma07_19880* and *Ma07_19890* could be (partly) involved in distinct, tissue-specific aspects of secondary mebabolite biosynthesis. The functional categorisation and its expression domains make *Ma07_19880* a good candidate to encode a proanthocyanidin biosynthesis regulator in developing banana fruits (pulp and peel), and thus an excellent target for genetic manipulation of proanthocyanidin content, known to influence senescence of banana fruit [46], to maintain the freshness of harvested banana fruit.

## Conclusions

The present genome-wide identification, chromosomal organisation, functional classification and expression analyses of *M. acuminata MYB* genes provide a first step towards cloning and functional dissection to decode the role of *MYB* genes in this economically interesting species. Further, knowledge of selected *MYB* genes can be biotechnologically utilized for improvement of fruit quality, yield, disease resistance, tolerance to biotic and abiotic stresses, and the biosynthesis of pharmaceutical compounds.

## Authors’ contributions

AP and RS conceived and designed research. BP and RS conducted experiments. BP analyzed and visualized transcriptome data. RS and BW interpreted the data. RS wrote the manuscript. All authors read and approved the final manuscript.

## Acknowledgements

We are grateful to Melanie Kuhlmann for excellent technical assistance. We thank Nathanael Walker-Hale for language editing.

## Supporting information

**S1 Table. MYB domain consensus sequence of R2R3-MYBs, used for tBLASTn searches.**

**S2 Table. RNA-seq raw data used for expression analysis.**

**S1 File. Multi-FASTA file with *MaMYB* CDS sequences.**

**S2 File. Multi-FASTA file with *MaMYB* peptide sequences.**

**S3 File. *MaMYB* genes general feature format (GFF) file.** For use in genome viewers/browsers on the *M. accuminata* pseudochromosomes (musa_acuminata_v2_pseudochromosome.fna).

