## Supplementary Tables 1 and 2 for "The *R2R3-MYB* gene family in banana (*Musa acuminata*): genome-wide identification, classification and expression patterns"

### Supporting information

**S1 Table. MYB domain consensus sequence of R2R3-MYBs, used for tBLASTn searches.**

GxWxxxEDxxLxxxxxxxGxxxWxxxxxxxGLxRxxKSCRLRWxNYLxPxxxxGxxxxxExxxxxxLxxxxGNxWxxIAxxxPxRTxNxxKNxW

**S2 Table. RNA-seq raw data used for expression analysis.**

| **biolocial source** | **Sequence Read Archive IDs** |
| --- | --- |
| embryogenic | SRR8761343,SRR8761344,SRR8761345 |
| seedling | SRR7340404,SRR7340406 |
| root | SRR2984591,SRR2984594,SRR2984596,SRR2984598,SRR2984601, SRR4450942,SRR4450934,SRR4450941,SRR4450933,SRR4450944 |
| leaf | SRR6320573,SRR6320439,SRR6320438,SRR6320441,SRR6320440, SRR6320443,SRR5405140,SRR5405141 |
| young_leaf | SRR6320573,SRR6320439 |
| adult_leaf | SRR6320438,SRR6320441 |
| old_leaf | SRR6320440,SRR6320443 |
| pulp | SRR5405133,SRR6320444,SRR5405132,SRR6320447,SRR5405131, SRR6320446,SRR5405130,SRR6320531 |
| pulpS1 | SRR5405133,SRR6320444 |
| pulpS2 | SRR5405132,SRR6320447 |
| pulpS3 | SRR5405131,SRR6320446 |
| pulpS4 | SRR5405130,SRR6320531 |
| peel | SRR5405139,SRR5405138,SRR5405137,SRR6320442,SRR5405136, SRR6320445,SRR5405135,SRR5405134 |
| peelS1 | SRR5405139,SRR5405138 |
| peelS2 | SRR5405137,SRR6320442 |
| peelS3 | SRR5405136,SRR6320445 |
| peelS4 | SRR5405135,SRR5405134 |
